# Kinetics and Evolution-guided Design of High Fidelity CRISPR-Cas Endonucleases

**DOI:** 10.64898/2026.07.06.736881

**Authors:** Huihui Liu, Zhenyu Zhou, Luowei Yuan, Bin Pang, Kun Xi, Xinyu Li, Wenzhuo Ma, Rujuan Ti, Jinchu Liu, Nuo Chen, Yang Xu, Jianyu Yang, Yiping Yu, Yuchen Yang, Ruobing Ren, Arieh Warshel, Yong Lei, Lizhe Zhu

**Author notes:** Corresponding (R.R.); (A.W.); (Y.L.); (L.Z.). These authors contributed equally (Huihui Liu, Zhenyu Zhou, Luowei Yuan).

## Abstract

Finding high-fidelity CRISPR-Cas variants is critical for both the precision of *in vitro* DNA detection and the safety of *in vivo* gene-editing therapeutics. However, the large size of the Cas enzymes and the distinct selection criteria between its natural evolution and clinical practice lead to extensive experimental trials and limited success rate for de novo design, directed evolution and protein language model (PLM)-based filtering. Here, we present a physics-driven, PLM-assisted approach that utilizes atomistic molecular dynamics simulations and automated path searching to efficiently obtain the atom-level kinetic insights, including the transition state structures, for the conformational changes of R-loop formation that dictate the off-target DNA recognition and cleavage by Cas. We show that these kinetic insights can pinpoint the fidelity-diminishing protein residues during the early stage of target recognition. Integrated with PLM-filtering of the mutants on these sites, this approach identified high-fidelity SpyCas9 and FnCas12a variants (4.49-fold and 6.69-fold increase respectively from wild-type), surpassing previously reported counterparts at minimal cost of only a few wet-lab validations (cellular genome-editing and ribonucleoprotein cleavage assays respectively).

## Introduction

The CRISPR-Cas systems (clustered regularly interspaced short palindromic repeats, CRISPR-associated proteins) are potent and versatile platform for genome editing^1–3^. Its programmability and operational simplicity have enabled wide applications across both basic research and therapeutic development, including clinical therapies for genetic disorders^4,5^. However, CRISPR-based clinical approaches still face a number of challenges, especially the unintended cleavage that may manifest the endo-nucleolytic activity at genomic loci and bear a few mismatches to the guide RNA (gRNA). This off-target cleavage is particularly concerning for *in vivo* therapeutics, where unintended edits can cause adverse outcomes^6–8^.

Despite its importance, finding high-fidelity Cas proteins have been largely restrained by the size of the enzyme and the distinct selection criteria between natural evolution and clinical practice. The ∼1000 sites of Cas proteins for random or site-saturation mutagenesis have posed daunting challenges for de novo design^9^ and directed evolution that rely on high-throughput wet-lab screening^10–13^. Therefore, protein language models (PLM) have been introduced for virtual filtering of candidate sequences and the generation of recombinant high-fidelity candidate variants from natural Cas sequences to reduce the load of wet-lab trials^14^.

However, as the role of Cas systems in the natural evolution of bacteria is to recognize and cleave the invading virus DNA that could be a family member of the recorded ancestor sequence in the gRNA, natural Cas proteins can barely hold ultra-high fidelity (Fig. 1a), leading to limited success rate of PLM-based filtering at the expense of hundreds of functional assays^15^. Consequently, finding a new source of information orthogonal to natural evolution seems an inevitable step towards a higher success rate of the design of high-fidelity Cas variant.

**Fig. 1.**
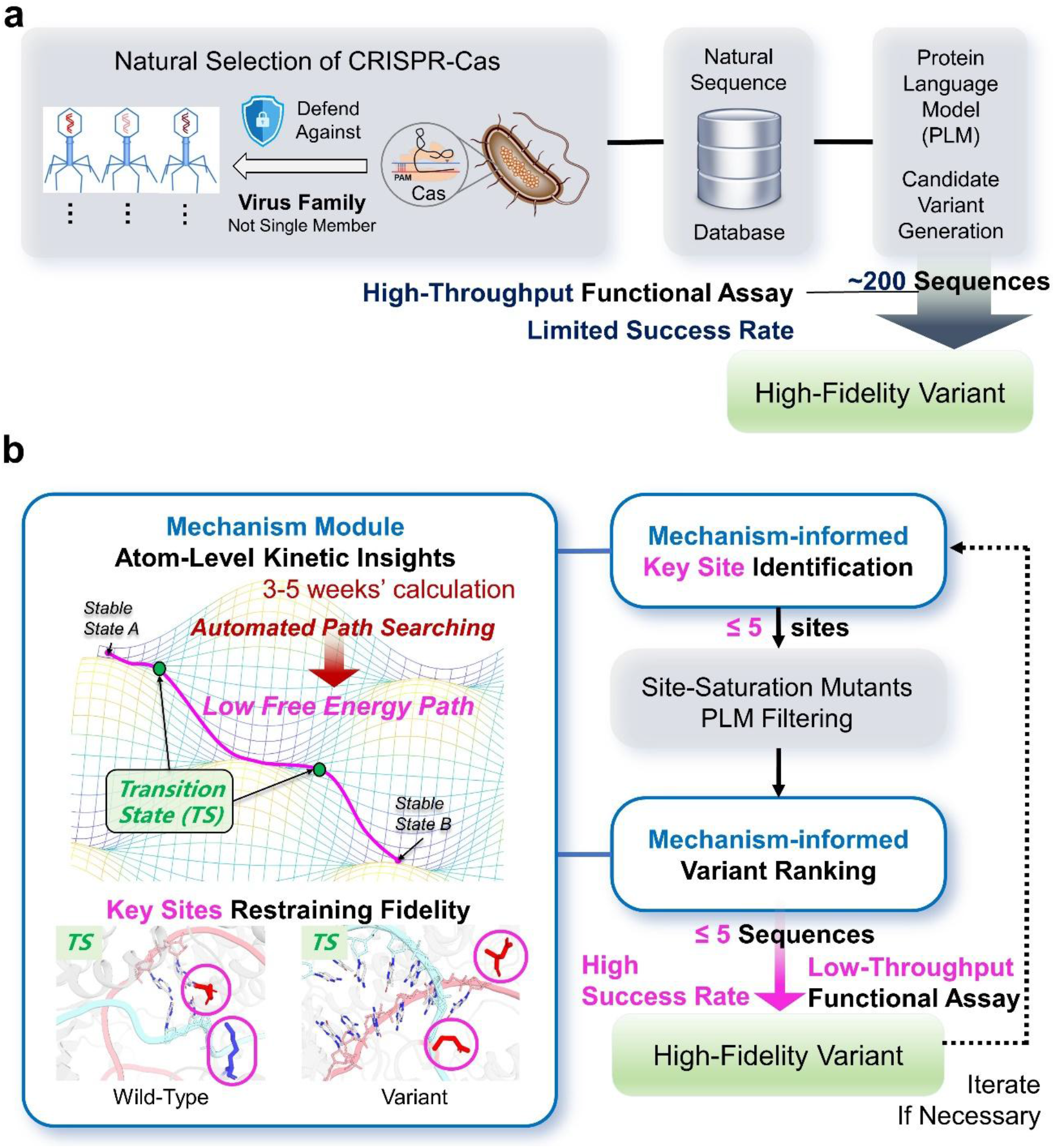
Comparison between protein language model (PLM)-based filtering of high-fidelity CRISPR-Cas variant and the mechanism-informed design in this work. **a.** Natural evolution selects Cas systems that defend the bacteria against virus families rather than a single family member, leading to the lack of high-fidelity of variants in the natural sequence database and therefore the limited success rate of the generation of fidelity-enhanced variants based on PLMs. **b.** Kinetic insights into the functional conformational changes of the Cas, if available, can pinpoint the critical enzyme residues that restrain fidelity, reduce the necessary throughput of functional assay and significantly increase the success rate for high-fidelity variant design.

Mechanistic insights into the catalytic cyle of Cas systems could be a complementary information source for this task. Mechanistic information could pinpoint key protein residues responsible for off-target cleavage and therefore significantly reduce the number of candidate sites for subsequent mutagenesis experiments (Fig. 1b). Nonetheless, obtaining the complete mechanism of a multi-domain complex enzyme itself has been a long-standing challenge.

The activity and specificity of an enzyme is defined by the overall rate of its catalysis on different substrates, which is governed by the rate-limiting (slowest) step with the highest kinetic barrier of its catalytic cycle. For CRISPR-Cas, it has been clear that this rate-limiting step is not the chemical reaction, but the conformational changes of substrate loading, i.e. the recognition of DNA by the Cas-gRNA complex, known as R-loop formation^16^. Therefore, resolving the free energy landscape underlying R-loop formation, especially the structure of the transition states (TS) that determine the rate of these conformational changes, is a prerequisite for precise tuning of Cas fidelity. However, transition states are too short-lived to be captured by structural biology experiments and too rare to observe in brute-force molecular dynamics (MD) simulations suffering from its slow computational speed brought by the expensive force calculations. Meanwhile, accelerating the search of transition states through algorithm innovation will inevitably be hindered by the curse of dimensionality: the acceleration itself requires prior knowledge of the location of transition states in the high dimensional space of the complex conformational changes, which is against the original goal of the search.

Here, we utilize our recently matured Travelling-salesman based Automated Path Searching (TAPS) algorithm^17–19^ to alleviate the curse of dimensionality and efficiently sample the low free energy path, covering all (meta)stable and transition state conformations, along the early stage of R-loop formation for two of the mostly widely used Cas tools (SpyCas9 and FnCas12a). Coupling this mechanism-aimed strategy with PLM-based filtering that ensures the thermal stability, folding integrity and other evolution-encoded properties of the candidate mutant sequences, we are able to identify SpyCas9 and FnCas12a with ultra-high fidelity (off-target cleavage rate 20-25% of wild-type for N4-Cas9 and 10-15% for Cas12a D194K-K1065D variant), surpassing previously reported counterparts at minimal cost of mutating only a few fidelity-diminishing residues and low-throughput functional assay validations.

## Results

### Target recognition for Cas9 seed region

As the most widely adopted gene-editing tool, SpyCas9 has been extensively studied using methods including structural determination, biochemical experiments and computational simulations^20–31^. Directed evolution has screened high-fidelity variants such as Sniper-Cas9^13^ and HiFi-Cas9^11^. Recently, PLMs succeeded in generating high-fidelity Cas9-like OpenCRISPR-1 from millions of candidates^15^. However, the selection and identification of high-fidelity variants in these methods heavily rely on high-throughput wet-lab experiments.

Mechanism-based rational design can in principle reduce the experimental load. Structure-guided engineering, for example, has found high-fidelity variants such as SpCas9-HF1^32^, eSpCas9^33^ and LZ3-Cas9^12^. These stable state structures, though offering a solid foundation for understanding its catalytic cycle, are not informative of the mechanism underlying the critical kinetic rates of the enzyme conformational changes, leaving systematic rational design beyond reach.

After the initial binding to the protospacer-adjacent motif (PAM), Cas9 undergoes REC-domain rearrangements that bend, distort, and partially unwind duplex DNA, enabling gRNA invasion and R-loop formation (gRNA-tDNA hybridization with formation of 20-nt length base pairs). Unlike the distal region (nt 11-20), the seed region (nt 1-10) dictates the initial target interrogation and serves as the primary determinant of specificity. However, improving the fidelity of the seed region (nt 1–10) is particularly challenging due to the hardly captured transient conformations that discriminate between on-target and mismatched sequences during this early phase.

Through the physics-driven TAPS approach with minimal requirement on guess inputs^17^, we resolved the low free energy path (LFEP) for the early stages of SpyCas9 R-loop formation in the seed region (Fig. 2a). The free energy distribution along the LFEP (Fig. 2b and 2c) reveals eight stages (Phase A-H) of this process: (A) firstly dsDNA starts to move towards REC3; (B) gRNA then initials pairing with tDNA; (C) recognition at nt 1-5 is accomplished; (D) Y450 forms contact with tDNA6; (E) separation of the HNH–REC3 interface occurs allowing the release of the 3′ tail of gRNA; (F) the DNA duplex starts to bend; (G) the DNA duplex starts unwinding from 7th position; (H) finally, R-loop forms at nt 7-10. (Supplementary Fig. 1) The heights of the 10 transition states (TS1-TS10) during these phases vary from 3 to 8.5 kcal/mol.

**Fig. 2.**
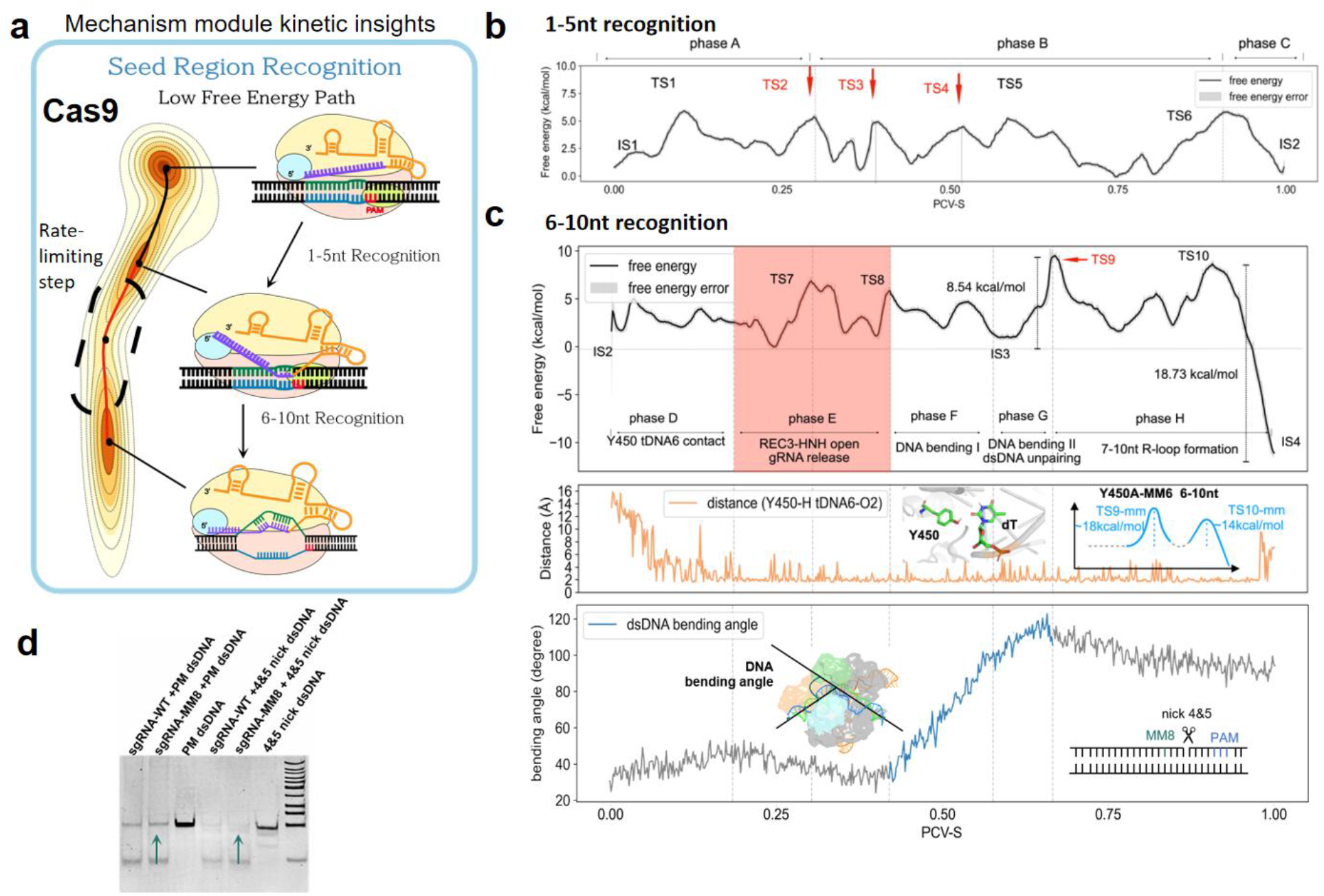
Seed region target recognition in SpyCas9. **a**. Schematic representation of seed region target recognition in SpyCas9. A low free-energy path for target DNA recognition in the seed region identifies residues critical for modulating mismatch formation and off-target cleavage, and informs biophysical descriptors used in the design. **b**. Free-energy profile showing the low free energy path for 1-5 nt recognition. **c**. Kinetic mechanism of target recognition at 6-10 nt in the seed region of SpyCas9. (Upper panel) Free-energy profile for 6-10 nt recognition; (Middle panel) Distances between Y450 and the 6th tDNA position. The energy schematic “Y450A-mm6” denotes the Y450A mutation with a gRNA-tDNA mismatch at position 6, and in this diagram the energy barriers for TS9-mm6 (∼18 kcal/mol) and TS10-mm6 (∼14 kcal/mol) are both markedly elevated compared to the WT on-target TS9 (∼8.5 kcal/mol) and TS10 (∼7 kcal/mol) in the 6-10 nt region; (Lower panel) DNA bending angles during the pairing process. **d**. Overlaid gel electrophoresis results illustrate that introducing a nick increases DNA bending flexibility, leading to higher off-target cleavage when a mismatch is present at position 8 (nick4&5DNA+MM8).

The rate-limiting step for seed recognition lies in Phase G, during which the DNA duplex bending angle reaches its maximum at the top barrier TS9 and DNA duplex unwinding starts from 7th position (Fig. 2c). To further verify the key role of such bending, we conducted cleavage assays of nicked dsDNA substrates with wild-type (WT) SpyCas9. These experiments show that the presence of a nick within the seed region significantly enhances SpyCas9’s tolerance for mismatches, pointing to the critical role of DNA bending rigidity in mitigating off-target cleavage in the seed region (Fig. 2d and Supplementary Fig. 7). In addition, prior to gRNA release (Phase E) and the dsDNA bending (Phase F & G), Y450 was found to form a stable tripartite structure with the 6th tDNA/gRNA base (Phase D, Fig. 2c) that persists until the seed-recognized state IS4 is fully reached. This indicates a positive role of Y450 in facilitating DNA bending and the recognition of both on-target and off-target sequences at nt 6, consistent with the previous experimental finding on the Y450A mutant^24^ and our additional simulations (Supplementary Fig. 3).

### Design of high-fidelity Cas9 variants

Further structural inspection (Fig. 2c) shows that the DNA bending starts after the release of the 3′ tail of gRNA (Phase E, Fig. 2c and Supplementary Fig. 4), which is constrained by five positively charged residues located on the L1 loop and bridge helix (BH) (R63, R66, K772, Q774, K775 in Fig. 3a). These residues limit the mobility of gRNA by long side chains and electrostatic interactions, thereby ensuring its proximity to tDNA for subsequent base-pairing (Supplementary Fig. 8). We therefore hypothesized that weakening such confinement could reduce the stability of mismatched gRNA-tDNA intermediates, thereby lowering off-target cleavage in the seed region.

**Fig. 3.**
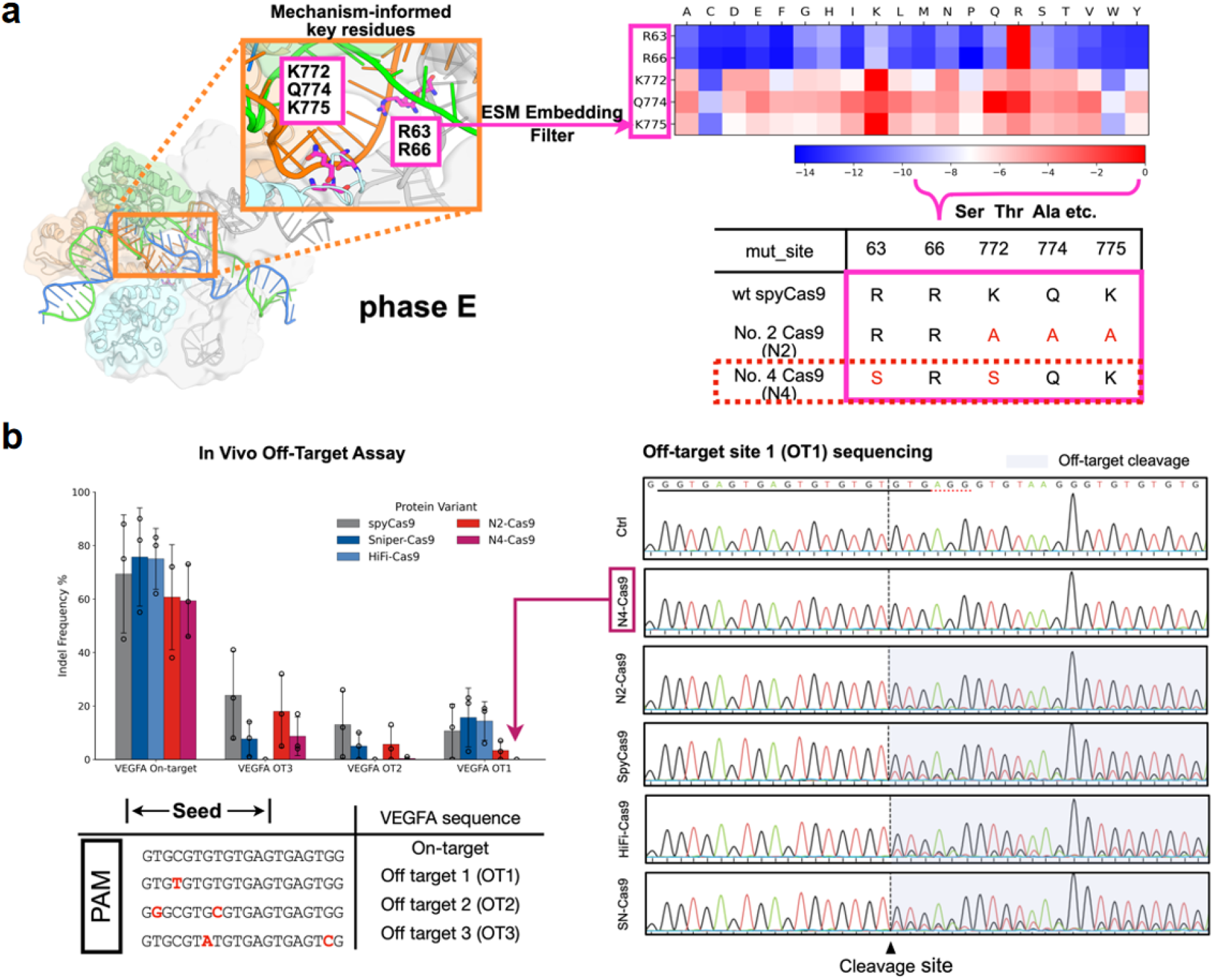
Mechanism-inspired design of Cas9 fidelity-enhancing mutants and functional validation. **a**, (Left panel) Structural diagram showing the position of five key positively charged long-side chain residues (R63, R66, K772, Q774, K775) within the separation process of the REC3 and HNH domains (phase E) and (right panel) heatmap of ESM embedding scores for all 20 substitutions at R63, R66, K772, Q774, K775 (rows = sites; columns = A-Y). Computed with esm2_t36_3B_UR50D via ESM-Scan. Color: blue = more negative (more deleterious). **b**, The left panel compares the intracellular editing performance of the newly designed N2-Cas9 and N4-Cas9 mutants with three widely used Cas9 nucleases (SpyCas9, HiFi-Cas9, Sniper-Cas9) at the *VEGFA* gene locus. Bars represent mean +/- standard deviation. (Right panel) Representative Sanger sequencing chromatograms for the OT1 site. The dashed line indicates the predicted cleavage site. The presence of overlapping peaks downstream of the cleavage site is indicative of heterogeneous indels resulting from nuclease activity.

All possible single-site substitutions at these positions were further scored with the protein language-model metric (ESM2)^34^, to safeguard the structural integrity and stability of the variants. Top 50% ESM embedding scores were retained for further design. Notably, ESM2 filtering alone seems inadequate for identifying these sites: R63 and R66 are highly conserved across Cas9 orthologs and flagged by ESM2 as evolutionarily “forbidden” for engineering.

The combinatorial variants are hypothesized to (i) leave the global Cas9 architecture intact, (ii) maintain essential contacts required for on-target cleavage, (iii) relax the electrostatic cage surrounding the gRNA, reducing the stability of mismatched gRNA-tDNA intermediates and (iv) consequently diminish off-target cleavage in the seed region. With such strategy, we obtained two variants N2(772A,774A,775A) and N4 (61S,770S) for downstream biochemical and cellular validation.

We benchmarked two engineered variants (N2-Cas9, N4-Cas9) against three established nucleases (SpyCas9, HiFi-Cas9, Sniper-Cas9) at the *VEGFA* locus. While all enzymes achieved robust on-target editing, their performance on off-target editing diverged across the three known off-target sites (OT1–OT3) (Fig. 3b). Relative to SpyCas9, both N2-Cas9 and N4-Cas9 showed markedly reduced indel frequencies at all off-targets (OT1-OT3). HiFi-Cas9 and Sniper-Cas9 also lowered off-target rates at OT2 and OT3, with HiFi-Cas9 exhibiting the strongest reduction at these two sites among all tested variants. At OT1, N4-Cas9 displayed the highest specificity, with indels at 0.0 ± 0.0%. SpyCas9 produced prominent mixed peaks initiating at the cleavage site (triangle), consistent with low fidelity joining after off-target cleavage. In contrast, N4-Cas9 yielded a near-baseline trace comparable to the no-edit control, providing sequence-level evidence of minimal off-target activity at OT1.

Base-resolution mutation rate profile (Supplementary Fig. 9) quantitatively confirmed these findings, showing widespread high-frequency mutation rates across OT1 in SpyCas9-treated cells versus minimal background-level mutation rates in N4-Cas9 samples, calculated from Sanger peak values. Extended indel spectrum analysis (Supplementary Fig. 10) demonstrated that SpyCas9 generated diverse indels at all off-target sites (OT1-OT3), while N4-Cas9 produced dramatically simplified profiles dominated by the WT sequence. The sanger sequencing chromatograms of OT2 and OT3 (Supplementary Fig. 11) also indicated the off-target reduction of N4-Cas9.

### Key residues for Cas12a seed fidelity

CRISPR-Cas12a is among widely used CRISPR-Cas systems for high-precision *in vitro* DNA detection^35^. After the R-loop formation, Cas12a cleaves not only the designated sites of targeted DNA sequence (*cis*-cleavage), but also single strand DNA (ssDNA) non-specifically (*trans*-cleavage) at low temperature^36^. The *trans*-cleavage offers a natural mechanism to amplify the signal of target recognition, when ssDNAs are labeled by fluorophore for reporting the *cis*-cleavage, presenting Cas12a a practical tool for pathogen detection and genotyping^37,38^. However, the precision of such DNA detection remains limited due to the off-target cleavage by Cas12a.

The design and screening of high-fidelity CRISPR-Cas12a faces the same heavy wet-experiments loads as CRISPR-Cas9 using directed evolution and PLM methods. Structure-guided engineering has generated high-fidelity AsCas12a (HyperFi-As) which lowered tolerance to mismatch at the position of the PAM-distal region^39^. However, the existing structures are mere stable snapshots without insights into the kinetics of its catalytic cycle and therefore barely informative of the reason behind the off-target recognition at the seed region—the most critical region that determine specificity^40–44^.

To find high-fidelity CRISPR-Cas12a variant at minimal cost, we adopted the physics-driven PLM-assisted workflow as described in Fig. 1b and 4a. We first located the LFEP underlying the seed region (nt 1-8) recognition of the target DNA by the WT FnCas12a. As shown in the free energy surface along the located LFEP (Fig. 4c), this process is featured by a top energy barrier of 8.7 kcal/mol, ultimately resulting in a stable state IS5 with lower energy than the initial state. Our simulation results indicate that the 8-bp structure is a stable intermediate state, which is consistent with the crystallographic structures^42^. Throughout the entire pairing process, the conformational changes of the protein were relatively small (Supplementary Fig. 12). The rate-limiting step is the first step corresponding to the proximity of the bases at positions 3 and 6. Structural analysis shows that during the transition of IS1 to IS2, the K1065 side chain changes from interacting with tDNA-6 phosphate to interacting with tDNA-5 phosphate (Fig. 4d and Supplementary Fig. 13). Subsequently, K1065 maintained its interaction with tDNA-5 phosphate, stabilizing the phosphate backbone (Supplementary Fig. 13).

**Fig. 4.**
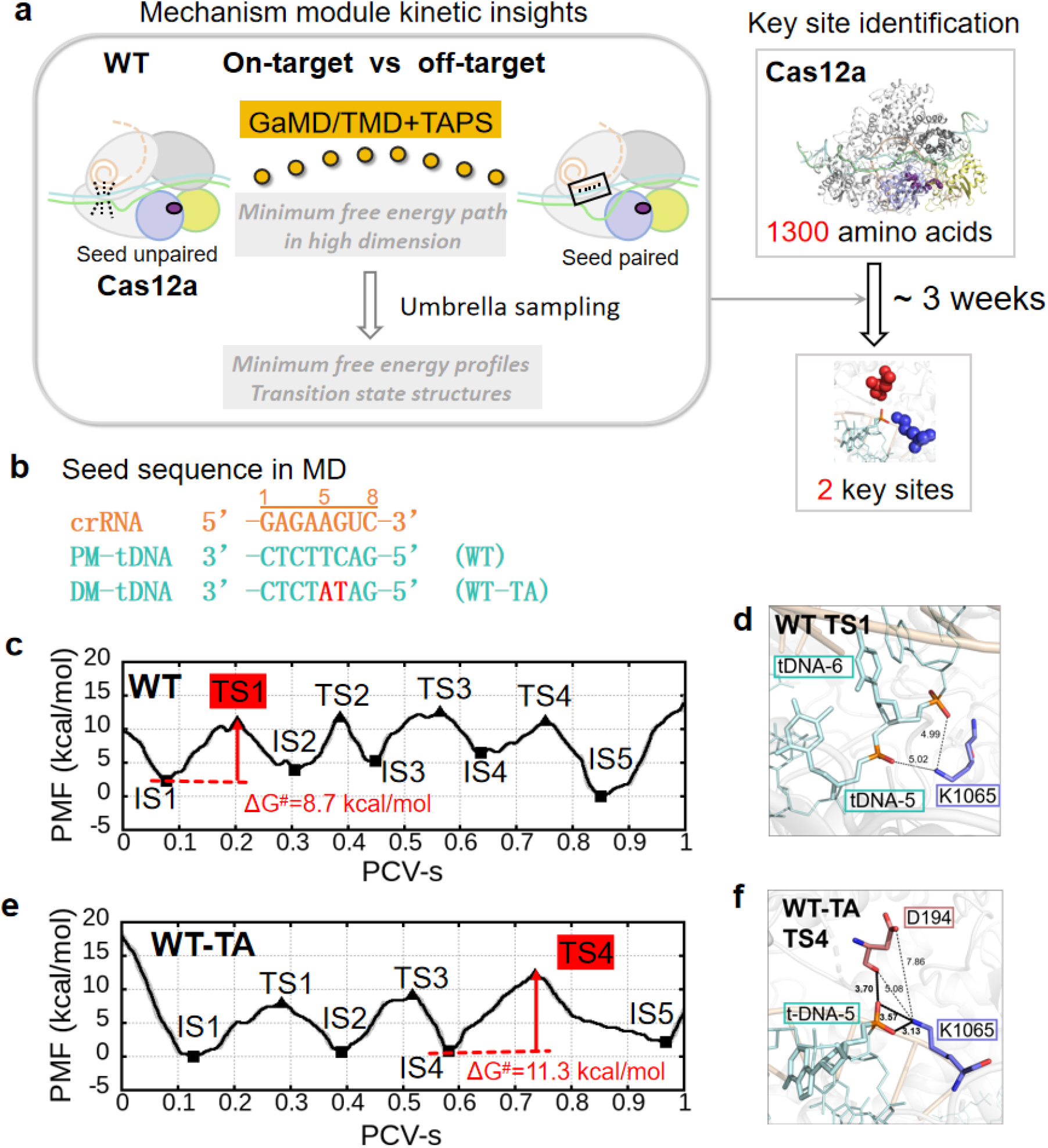
Key site identification for CRISPR-Cas12a. **a**, Workflow for low free energy path searching and key sites identification. **b**, Perfect match and designed mismatched DNA sequences used in MD simulations. **c**, The potential of mean force profile for WT FnCas12a with perfect match DNA sequence and the rate-limiting step structure (**d**, WT TS1). **e**, The potential of mean force profile for WT FnCas12a with TA mismatched DNA sequence and the rate-limited step structure (**f**, WT-TA TS4).

FnCas12a has been reported with high selectivity at positions 1-6 except position 5^45^. In order to investigate the reasons for off-target recognition at position 5, we designed a double mismatched sequence at positions 5 and 6 (Fig. 4b, Supplementary Table 2) and obtained the low free energy path for mismatched sequence pairing (WT-TA). The free energy profile for WT-TA is shown in Fig. 4e. Due to the 5 and 6 dislocation pairing, the free energy barrier of the last transition state (TS4) is the highest (11.3 kcal/mol), corresponding to the pairing of sites 3 and 4. Meanwhile, in the IS4 state and afterwards, K1065 and D194 are getting closer, forming a pincer like structure that stabilizes the phosphate backbone (Fig. 4f and Supplementary Fig. 14). Due to the stabilizing effect of D194 (Fig. 4f), the energy barrier increase for mismatch pairing is only 2.6 kcal/mol compared to correct pairing, consistent with the high off-target rate at nt 5.

### K1065E reduces *cis*-cleavage off-target

Our transition-state analysis identified K1065 as a critical regulator of seed-region base pairing in WT FnCas12a (Fig. 4d). Comparative analysis of on-target and mismatched pairing further indicated that D194 cooperates in stabilizing the phosphate backbone of the tDNA (Fig. 4f). Guided by this mechanistic insight, we first focused on engineering the residue K1065 (Fig. 5a).

**Fig. 5.**
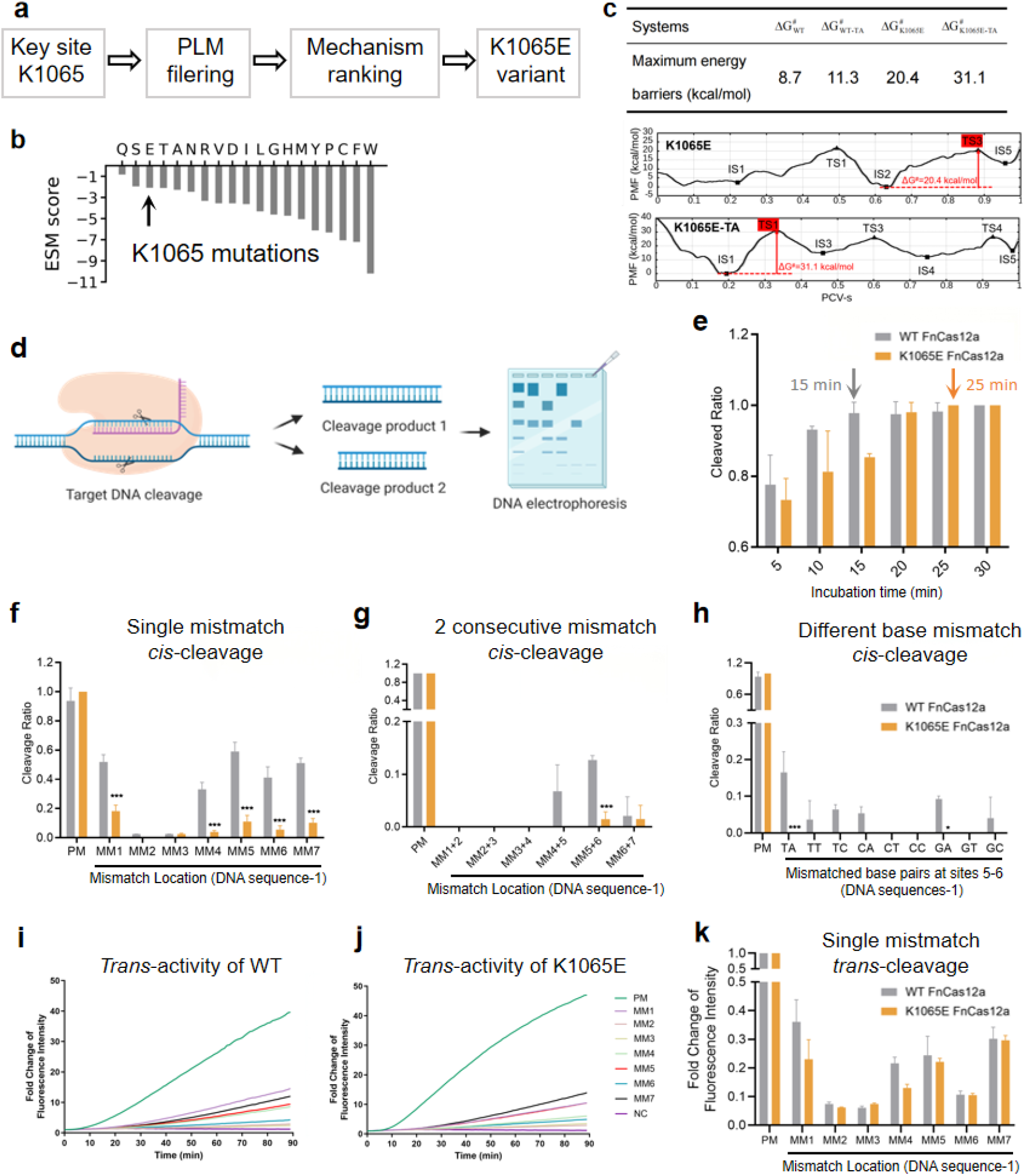
Cas12a K1065E mutant decreased off-target effect of sites 1-7 in *cis*-cleavage and increased *trans*-cleavage off-target discrimination by 66% at most. a, Workflow for Cas12a K1065E mutant design. **b,** ESM score given by ESM2 model. **c,** The potential of mean force profiles of base-pairing process for K1065E mutant. **d,** Experimental workflow for *in vitro* cleavage assays. **e,** Time-dependent cleavage kinetics of WT FnCas12a and the K1065E variant under optimized reaction conditions. *Cis*-cleavage activity in the presence of single-site mismatches (**f**) and dual consecutive mismatches within the seed region (**g**). **h**, Systematic base substitutions at positions 5 and 6 to assess sequence-composition effects. **i-k**, *trans*-cleavage activity triggered by target substrates containing single-site mismatches at different seed positions. Three independent target sequences were evaluated; results for sequence 1 are shown, with data for sequences 2 and 3 provided in Supplementary Fig. 19-20.

To prevent the stable electrostatic interactions between K1065 and tDNA, we designed K1065E variant which was ranked third by the ESM2 model^34^ (ESM2_t36_3B_UR50D) (Fig. 5b), supporting the tolerance of this mutation. To evaluate its impact on mismatch discrimination, we computed the low free-energy paths for K1065E during recognition of matched and mismatched DNA substrates (Fig. 5c). In contrast to WT, substitution of lysine with glutamate abolished stabilization of the tDNA phosphate backbone at position 5 (Supplementary Fig. 16), leading to an elevated free-energy barrier along the pairing pathway (Fig. 5c). In the presence of mismatches, the combined effects of electrostatic repulsion and disrupted backbone stabilization further increased crRNA-tDNA separation, hindering formation of the first base pair (Supplementary Fig. 16b). Consequently, the initial transition state became the dominant energetic barrier during the off-target recognition by K1065E. Relative to matched pairing, K1065E increased the activation barrier for mismatched substrate recognition by 10.7 kcal/mol (Fig. 5c), presenting a substantial reduction of off-target recognition and cleavage.

The performance of the K1065E variant was further evaluated using optimized *in vitro* ribonucleoprotein (RNP) cleavage assays (Fig. 5d-e and Supplementary Fig. 17). Compared with WT FnCas12a, the mutant showed markedly reduced cleavage when single or dual consecutive mismatches were introduced at PAM-proximal positions 1-7 (Fig. 5f-g and Supplementary Fig. 18a-b), consistent with suppression of seed-region off-target activity. To exclude potential bias related to purine-pyrimidine composition, we systematically substituted nucleobases at positions 5 and 6 (Supplementary Fig. 18c) and quantified cleavage efficiencies across all combinations. WT FnCas12a exhibited reduced specificity when one or both mismatched bases were adenine or thymine. In contrast, off-target cleavage by K1065E was effectively eliminated across all tested substitutions (Fig. 5h and Supplementary Fig. 18c). Similar improvements in specificity were also observed at additional previously reported target sites (Supplementary Fig. 19 and 20). These evidences all indicate that the enhanced fidelity by the K1065E substitution is largely sequence-independent rather than base-preferential.

Given the widespread use of Cas12a in nucleic acid detection, we next examined collateral *trans*-cleavage activity. RNP-based assays demonstrated that K1065E exhibited substantially reduced *trans*-cleavage activation in the presence of mismatches at positions 1 and 4 (Fig. 5i-k), whereas activity was comparable to WT at other mismatch positions. The transition from target-dependent *cis*-cleavage to collateral *trans*-cleavage was attenuated in the K1065E variant (Supplementary Fig. 21), suggesting that altered seed-region pairing kinetics modulate downstream conformational activation.

### Enhancing *trans*-cleavage discrimination

Comparative analysis of on-target and mismatched pairing in WT FnCas12a revealed that D194, together with K1065, contributes to stabilization of the tDNA phosphate backbone. We therefore hypothesized that coordinated modulation of these two residues could further improve mismatch discrimination. To disrupt the electrostatic stabilization of tDNA while minimizing structural perturbation to the protein scaffold, we engineered a charge-swapped double mutant, D194K-K1065D. ESM2 prediction^34^ (ESM2_t36_3B_UR50D) indicates that the D194K substitution is tolerable (Fig. 6a).

**Fig. 6.**
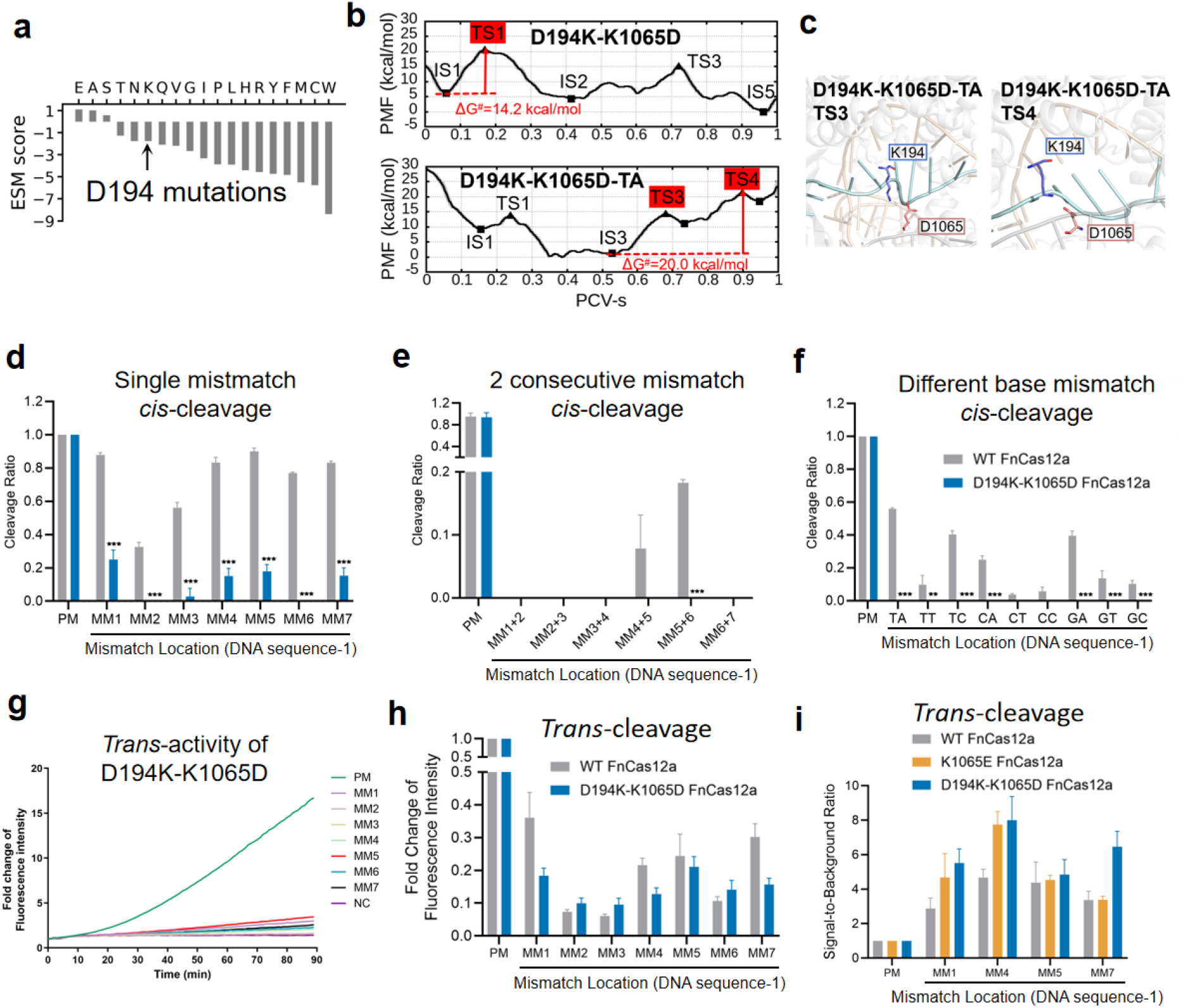
Cas12a D194K-K1065E mutant decreased off-target effect of sites 1-7 in *cis*-cleavage and increased *trans*-cleavage off-target discrimination by 92% at most. a, Predicted thermostability scores of the D194K-K1065D variant generated by the ESM2 model. **b,** Free-energy profiles (potential of mean force) for matched and mismatched DNA recognition by the D194K-K1065D variant. **c,** Representative transition-state structures along the mismatch recognition pathway, highlighting destabilization of seed-region pairing. *Cis*-cleavage activity in the presence of single-site mismatches **(d)** and dual consecutive mismatches **(e)** within the seed region. **f,** Systematic base substitutions at positions 5 and 6 to assess sequence-composition effects. **g-i,** *trans*-cleavage activity triggered by substrates containing single-site mismatches across seed positions 1-7. Three independent target sequences were examined; results for sequence 1 are shown, with data for sequences 2 and 3 provided in Supplementary Fig. 24 and 25. Maximum enhancement in mismatch discrimination reached 92% relative to WT FnCas12a.

To assess the energetic consequences of this design, we located the low free-energy path for D194K-K1065D during recognition of matched and mismatched substrates. The free energy landscape along the path is shown in Fig. 6b. For matched DNA, loss of stabilizing interaction between residue 1065 and the tDNA backbone increased the highest activation barrier to 14.2 kcal/mol (Fig. 6b), consistent with moderated pairing kinetics. For mismatched substrates, the base-pairing trajectory became trapped in intermediate state IS3 due to combined destabilization at positions 5 and 6 and loss of transition-state stabilization previously mediated by K1065 (Fig. 6c and Supplementary Fig. 22). As a result, the activation barrier exceeded 20 kcal/mol, markedly reducing the probability of off-target pairing.

We next validated these predictions experimentally. To maintain consistency with the K1065E analysis, cleavage reactions were performed above optimized condition. *In vitro* digestion assays demonstrated that *cis*-cleavage activity toward single and dual consecutive mismatches (sequence 1) was reduced by at least 72% relative to WT (Fig. 6d-f and Supplementary Fig. 23). Similar improvements in specificity were observed across additional published target sequences (Supplementary Fig. 24 and 25), supporting the robustness of the design.

Given the importance of collateral activity for diagnostic applications, we further evaluated *trans*-cleavage using RNP assays. The D194K-K1065D variant exhibited substantially reduced *trans*-cleavage activation in the presence of mismatches at positions 1, 4 and 7 (Fig. 6g-i). Except at sites where WT Cas12a already displayed intrinsically low off-target activity (positions 2, 3 and 6), *trans*-cleavage mismatch discrimination was enhanced across seed positions 1-7, reaching up to 92% improvement relative to WT (Fig. 6i).

Together, these results demonstrate that coordinated electrostatic reprogramming at residues 194 and 1065 synergistically reshapes the kinetic landscape of seed pairing, yielding a high-fidelity FnCas12a variant with markedly improved *trans*-cleavage specificity.

## Discussion

Rational enzyme design has been regarded as a valuable approach for guiding and minimizing downstream wet-lab trials, since the maturation of quantum mechanics/molecular mechanics (QM/MM) for dissecting the step of chemical reaction^46,47^. However, systematic rational design seldom became feasible due to the lack of kinetic insights into the complex conformational changes before and after the chemical reactions. The present work demonstrates that, such kinetic insights, once routinely obtainable by automated path-searching methods, can pinpoint critical residues limiting the enzyme activity and selectivity. Coupling the kinetic insights with PLM that prioritize the variants on the identified sites, we are able to enhance the fidelity of CRISPR-Cas systems at remarkably low cost, which is beyond the reach of coevolution-based approaches alone. Further coupling of this general kinetics-based PLM-assisted workflow with the QM/MM approach will secure a promising strategy able to systematically predict all transition state structures during the catalytic cycle of arbitrary enzymes for rational engineering.

## Methods

### Simulation system preparation

The initial structures of SpyCas9 in complex with single-guide RNA (sgRNA) and double-stranded DNA (dsDNA) were obtained from the Protein Data Bank (PDB IDs: 7S36, 7S38, 7Z4E, 7Z4D). Missing residues in DNA and protein loops were modeled using MODELLER^48–50^. Mutant structures (Y450A-OT and Y450A-MM6) were generated using PyMOL^51^ mutagenesis based on PDB structures. The simulation systems were constructed in a TIP3P solvent box at 310 K and pH 7.5, with 250 mM KCl and 5 mM MgCl₂ to mimic physiological conditions (Supplementary Table 3).

We used the Cryo-EM structure (PDB ID: 6GTC) as a template to construct a FnCas12a-crRNA-DNA ternary complex structure where crRNA and tDNA were not paired. The missing Cas12a amino acids were completed using MODELLER^48,49^, while crRNA and DNA were constructed using the MacroMoleculeBuilder (MMB) software^52^. The magnesium ions in the PDB structure were retained. The protonation states of titratable residues were determined based on the pKa predicted by PROPKA 3.1^53^ and pH 7.4. When constructing the system, we first dissolved the complex in water, and then added 150mM NaCl to neutralize the charge. The total number of atoms was 218,242. The initial structures for K1065E mutant and mismatch sequences were obtained using WT perfect match sequence structure as template by MODELLER^48,49^.

### Molecular dynamics (MD) simulations

MD simulations were performed using GROMACS 2019.4 or 2021.4^54^. AMBER force field was selected as the force field. Among them, ff14SB was used for protein force field^55^, OL3 was used for RNA^56^, OL15 was used for DNA^57^ and TIP3P model was used for water^58^. Energy minimization was conducted 10,000 steps of steepest descent followed by the conjugate gradient algorithm to remove steric clashes. After minimization, the system was equilibrated in two stages: NVT ensemble at 310 K for 1 ns to equilibrate the solvent and NPT ensemble for 1 ns at 1 atm. Production MD simulations used a 2-fs time step with periodic boundary conditions. Long-range electrostatic interactions were treated using the Particle Mesh Ewald (PME) method^59,60^, while short-range electrostatics and van der Waals interactions were handled with a 10 Å cutoff for Cas9 and a 12 Å cutoff for Cas12a. The LINCS algorithm^61^ was applied to constrain all bonds involving hydrogen atoms.

### Initial path generation

For CRISPR-Cas9, to generate the initial transition path of WT-OT 1-5 nt and WT-OT 6-10 nt before optimization, targeted MD (TMD) simulations^62,63^ were conducted using GROMACS 2019.4^54^ and PLUMED 2.5.3^64,65^. The reference structures used in TMD were: initial conformation (PDB ID: 7S36, equilibrated after 100 ns MD simulation), 3 nt structure (PDB ID: 7S38), 5 nt structure (PDB ID: 7Z4C), 8 nt structure (PDB ID: 7Z4E), 12 nt structure (PDB ID: 7Z4G). The alignment atoms were selected from Cα atoms of residues with minimal secondary structure variations (helix regions) between consecutive structures. The biased atoms included heavy atoms in regions exhibiting significant secondary structure changes and the phosphate backbone and sugar ring atoms in nucleotides undergoing conformational transitions. The biasing spring constant k(t) was applied as follows: (a) to 500 kJ/(mol·nm²) from 0 ns to 5 ns; (b) 500 to 500,000 kJ/(mol·nm²) from 1 ns to 10 ns. Initial transition path of Y450A-OT and Y450A-MM6 were constructed based on optimized Y450A-MM6 trajectory. To extract the initial path for later optimization, the TMD trajectory was sliced at 1.55 Å intervals, illustrated in Supplementary Table 4.

The initial pathway for the WT Cas12a protein to recognize correctly paired target DNA was extracted from conventional MD simulations and Gaussian accelerated MD (GaMD) simulations^27,28,66^. The molecular simulation software used was AMBER18^67^. Starting from the initial structure above, we first used the steepest descent algorithm and conjugate gradient algorithm to minimize system energy. Then water and ions were relaxed in the NVT ensemble, following by slowly releasing of solute constraints in the NPT ensemble. Finally, 100 ns unconstrained equilibrium simulation were performed. Temperature was 310K controlled by Langevin thermostat and pressure was 1 standard atmospheric pressure maintained by Berendsen barostat. Then we run a 108.8 ns GaMD pre simulation, followed by an 800 ns GaMD. The dihedral potential energy and total potential energy are both biased to enhance the acceleration. The initial paths for mismatched sequences and K1065E mutant were obtained by targeting the structures on the correct pairing process pathway of WT Cas12a obtained from GaMD step by step using TMD.

### Path optimization

The TAPS (Traveling-Salesman-Based Automated Path Searching) method was utilized to refine the initial transition path into the low free energy path (LFEP). The theoretical background and methodological details of TAPS have been previously documented in our published work^17–19,68^. Path optimization was performed using a custom Python script (https://github.com/liusong299/TAPS). The convergence of the optimized pathway was validated through 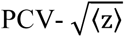 analysis, as illustrated in Supplementary Table 5 and Supplementary Fig. 26 and 27.

### Free energy calculations

To evaluate the free energy landscape along the LFEP and characterize key transition and intermediate states, we conducted umbrella sampling simulations^69^ using GROMACS^54^ in combination with PLUMED^64,65^. The free energy profiles were constructed along the PCV-s coordinate, providing insight into the conformational transitions along the LFEP.

For Cas9, sampling windows were defined at intervals of 0.25 PCV-s along the LFEP, with each window subjected to a harmonic biasing potential using a force constant of 600 kJ/(mol·nm²). Additionally, a harmonic wall potential with a force constant of 3,152,000 kJ/(mol·nm²) was applied at PCV-z = 0.0256 to confine sampling within 0.04 nm of the LFEP. Each window was simulated for a minimum of 4 ns, with the first 200 ps discarded as equilibration. The weighted histogram analysis method (WHAM)^70^ was employed to reconstruct the full free energy profile. The parameter details for umbrella sampling are summarized in Supplementary Table 6.

For Cas12a, the sampling was performed along the LFEP at intervals of 0.2 PCV-s. Each window run 4 ns production run and the last 2 ns was used for analysis by the WHAM method^70^.

### MD simulations analysis

In order to analyze the domain interface dynamics of Cas9, the minimum distance between the RuvC and HNH domains was calculated using GROMACS^54^ with a cutoff of 1.5 nm.

Hydrogen bonds between SpyCas9 and DNA were quantified using MDAnalysis (version 2.7.0) with donor-acceptor distance < 3.5 Å and angle > 120°.

The DNA bending angle was calculated by measuring the angle between two vectors, each defined by the center of mass of specific segments at either end of the double-stranded DNA. For each vector, two groups of nucleotides were used: the terminal 2 base pairs and the adjacent 3 base pairs (i.e., the penultimate region). The center of mass of each group was first determined, and then a vector was constructed from the center of mass of the penultimate region to that of the terminal region. The angle between these two vectors—corresponding to the two ends of the DNA duplex—was then computed to represent the overall DNA bending angle (Supplementary Fig. 5).

### ESM-based mutational scoring

Residue-likelihood evaluations were carried out with a custom Python workflow adapted from the open-source ESM-Scan toolkit ( https://github.com/xuebingwu/ESM-Scan.git)^34,71^. The pipeline loads the esm2_t36_3B_UR50D pre-trained language model, tokenizes each Cas9/Cas12a variant carrying a single-site substitution, and computes the log-likelihood ratio (mutant versus WT) at the mutated position. Scores are then normalized across all candidates, and substitutions that incur minimal likelihood loss - implying limited structural or functional perturbation - are retained for downstream design.

### Plasmid design and construction

The coding sequences of HiFi Cas9 and Sniper Cas9 were synthesized commercially (GENEWIZ) and cloned into PX459 (Addgene #48139) backbone between the AgeI and FseI restriction sites. The variants N2 Cas9 and N4 Cas9 were cloned by site-directed mutagenesis into PX459. Site-directed mutagenesis was employed to generate N2 and N4 Cas9 variants using PX459 as the template. The VEGFA sgRNA cassette was constructed by annealing complementary pairs of oligonucleotides and ligating the DNA fragments into PX459 and its variants vectors between BbsI restriction sites. Ligated plasmids were transformed into E. coli DH5α competent cells. Single colonies were selected on LB-ampicillin agar plates, followed by plasmid extraction and Sanger sequencing to verify insert sequences and reading frames. The fully matched dsDNA and nicked DNA substrates were prepared by annealing complementary oligonucleotide pairs in 1× Annealing Buffer for DNA Oligos (Beyotime, D0251) using the following thermal profile: 95 °C for 5 min followed by a linear ramp to 25 °C at -0.1 °C/cycle (700 cycles total duration).

For Cas12a, targeted DNA products complementary to crRNA (Genscript) were performed by annealing complementary pairs of oligonucleotides (Supplementary Table 2) on ProFlex 3 x 32-well PCR System (Thermo Fisher Scientific) and ligating the DNA fragments into pUC19 plasmid between the EcoRI and HindIII restriction sites. Then, the ligated plasmids were transformed into E. Coli. After selecting the individual E. Coli colonies, all the sequences of the recombinant plasmids were confirmed by sanger sequencing.

### Ribonucleoprotein cleavage experiments

For Cas9 cleavage assays, 100 nM Cas9 (Genscript, Z03469) was pre-incubated with 120 nM sgRNA (Synthesized at Genscript) in 1x Cas9 Nuclease Reaction Buffer (Genscript, Z03469) at 37℃ for 10 min to form ribonucleoprotein (RNP) complexes. Subsequently, 40 nM DNA substrates were added to the RNP mixture (final volume 20 μL), and reactions proceeded at 37°C for 20 min. Reactions were terminated by adding 1 μL RNase A (TransGen Biotech, GE101-01) and 1 μL Proteinase K (TransGen Biotech, GE201-01) with incubation at 37°C for 20 min. DNA products were mixed with 6× DNA Loading Dye (TransGen Biotech, GH101-01), resolved on 15% TBE polyacrylamide gels at 135 V for 75 min, stained with 1× GelStain (TransGen Biotech, GS101-01), and visualized using a TANON Mini Space 2000 imaging system.

For Cas12a, 2 μg of plasmids were linearized by incubation with 40 units of SspI-HF (New England Biolabs) in 1x rCutSmart buffer for 2 h at 37 ℃ in a total volume of 20 μL, following heat inactivation for 20 min at 65 ℃. For cleavage assays, FnCas12a (Genscript) in SEC buffer (20mM HEPES-KOH pH 7.5, 500mM KCl, 1mM DTT) was incubated with crRNA and MgCl_2_ (Beyotime) at 37℃ for 10 min to allow binary complex assembly, followed by adding linearized plasmids. The reaction for the cleavage assays was performed by adding 10 μM FnCas12a (in SEC buffer) 5μL, 10 μM crRNA (in H_2_O) 2 μL, 100 mM MgCl_2_ (DEPC-treated) 1 μL and linearized plasmids DNA 250 ng. The total volume was adjusted to 20 μL using water. The reaction was incubated for 30 min at 37℃. Reactions were stopped by adding 0.5 M EDTA (Beyotime) and 20 mg/ml Proteinase K (Beyotime) to final concentrations of 80 mM and 0.8 mg/ml, respectively, for 30 min at 37℃. After adding 6X DNA loading dye (Beyotime), the DNA products were resolved on 0.8% agarose gels stained with Gel-Red (Beyotime) and visualized on ChemiDoc Imaging System (Bio-Rad). Data were performed as mean ± SD. Statistical significance was analyzed by using Student’s unpaired t-test in GraphPad Prism 6 (Graph-Pad Software, Inc, California, USA). ∗P < 0.05, ∗∗P < 0.01 and ∗∗∗P < 0.001 were regarded as statistically significant.

### Cell culture and transfection

HEK293T cells (ATCC CRL-3216) were provided by American Type Culture Collection (ATCC), and cultured in Dulbecco’s modified Eagle’s medium (DMEM) containing with 10 % (v/v) fetal bovine serum (FBS) in the condition of 5 % CO_2_ at 37 °C. For cell transfection, HEK293T cells were seeded at a density of 1 × 10⁵ cells per well in a 24-well plate and allowed to adhere overnight. The following day, 1 μg of plasmid was transfected into the cells using jetOPTIMUS Transfection Reagent (Polyplus, Catalog #101000006) according to the manufacturer’s protocol. After 24 h of incubation, transfected cells were selected by treatment with 4 μg/mL puromycin for 48 h. The total transfection period was 72 h.

### PCR and indel frequencies quantification

After puro selection, the genomic DNA of the transfected cells were collected using Animal Tissue Direct PCR Kit (Yeasen, Catalog #10184ES50). Target and predicted off-target regions flanking the sgVEGFA-directed Cas9 variant binding sites were amplified by PCR using GoTaq Green Master Mix (Promega Catalog #PAM7122) with primers spanning ∼300 bp upstream and downstream of each locus. Amplified products were purified and subjected to Sanger sequencing. Sequencing traces were analyzed using the Synthego ICE CRISPR Analysis Tool (https://www.synthego.com) to quantify insertion/deletion (indel) frequencies and editing efficiency at both on- and off-target sites.

## Acknowledgements

This research was supported by grants by:

Warshel Institute for Computational Biology, School of medicine, The Chinese University of Hong Kong, Shenzhen, Guangdong, China (No. LGKCSDPT2025001).

Department of Bioinformatics, School of Medicine, The Chinese University of Hong Kong, Shenzhen, Guangdong, China.

Science, Technology and Innovation Commission of Shenzhen Municipality (No. RCYX20200714114645019) and National Natural Science Foundation of China (No. 31971179) to L.Z.

National Natural Science Foundation of China (No. 32170583), Guangdong Pearl River Talents Program (No. 2021QN02Y438), and Shenzhen Natural Science Foundation (No. JCYJ20220818103008017) to Y.L.

## Author Contributions

**H.L. and Z.Z.** designed the study, performed MD simulations, path optimization, data analysis, and protein design, and wrote the manuscript. **L.Yuan., N. C. and Y.X.** performed protein expression, *in vitro* nucleic acid cleavage assays, *in vivo* cleavage assays, and off-target sequencing analysis. **B.P.** carried out protein expression and purification. **X.L., W.M., R.T., J.Y., and Y.Yu** contributed to data analysis and manuscript revision. **K.X., J. L. and Y.Yang.** assisted in figure preparation and revision. **R.R.** revised the manuscript. **A.W.** supervised the project, analyzed the data, and revised the manuscript. **Y.L. and L.Z.** supervised the project, designed the experiments, acquired funding, and revised the manuscript. **H.L.**, **Z.Z.** and **L.Yuan** contributed equally to this work.

## Data Availability Statement

See Supplementary materials.

Correspondence and requests for materials should be addressed to Lizhe Zhu or Yong Lei.

