## Supplemental Figures and Tables for "Kinetics and Evolution-guided Design of High Fidelity CRISPR-Cas Endonucleases"

**This PDF file includes:**

Supplementary Figs 1 to 27

Supplementary Tables 1 to 9

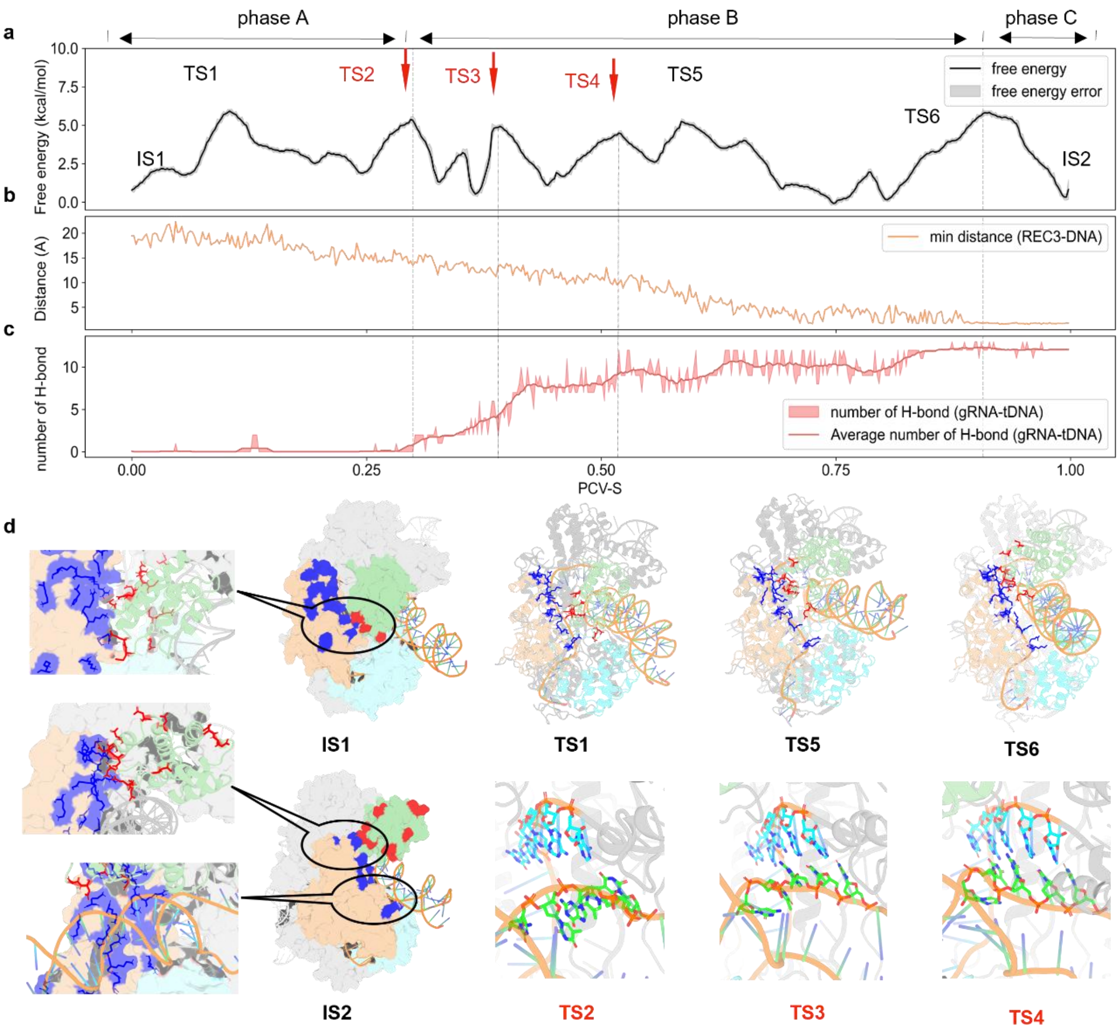


**Supplementary Fig. 1 | Kinetic Mechanism of Target Recognition at 1-5 nt by SpyCas9. a**. Free-energy profile showing the low free energy path for 1-5 nt recognition. **b**, Minimum distance between REC3 and DNA. **c**, Number of hydrogen bonds between gRNA and tDNA. **d**, Structural details of the electrostatic interactions between REC2, REC3, and dsDNA. Structural snapshots of TS1, TS5, and TS6, respectively, with the Rec2 domain colored green, Rec3 domain colored wheat, HNH domain colored cyan, positively charged residues in blue, and negatively charged residues in red. Structural snapshots of TS2, TS3, and TS4, respectively, with gRNA colored in cyan, tDNA in green.

During the initial Phase A, the negatively charged residues of REC2 begin sliding along the positive charge track on REC3, inducing a rotational movement of the REC2-DNA complex. As this rotation progresses (Phase B), REC2 pulls the DNA duplex toward REC3. During this transition, gRNA progressively form base-pairs with tDNA at nt 1-5. Upon completion of base-pairing, the DNA continues to move until REC2 flips above the DNA, anchoring the duplex onto the positive charge track on REC3 by overcoming the top barrier TS6 (5.83 kcal/mol) among the six (Phase C, IS2). Collectively, the six low barriers point to the key role of the sliding-encouraging charge distribution at the REC2/REC3 interface for facilitating dsDNA distortion and therefore accelerating gRNA-tDNA base-pairing at the first half of the seed region.

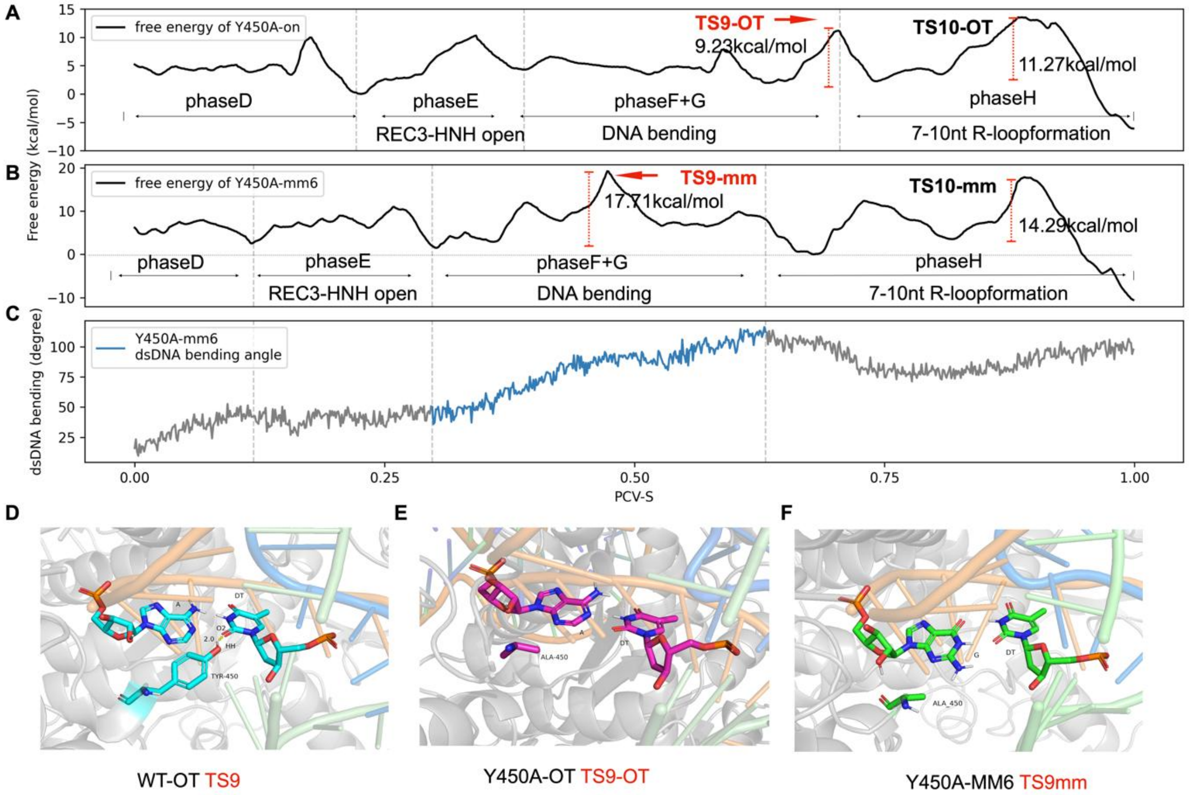


**Supplementary Fig. 2 | Y450 modulates DNA bending and mismatch tolerance in the seed region of SpyCas9. a-b**, Free-energy profiles of Y450A-PM (perfect match) and Y450A-mm6 (6th position mismatch). **c**, DNA bending angle in Y450A-mm6, showing TS9-mm occurs before maximum bending. **d-f**, Structural comparison of TS9 (wt-on), TS9-OT (Y450A-OT) and TS9-mm (Y450A-mm6).

In wild-type Cas9, Y450 forms a hydrogen bond with the 6th base of tDNA, stabilizing the local R-loop structure and facilitating DNA bending. To assess the role of Y450, three trajectories were compared: wt-on, Y450A-on (Y450A mutation with perfect match), and Y450A-mm6 (Y450A mutation with a mismatch at position 6). While key features such as REC3-HNH interface separation, DNA bending, and 7-10 nt hybridization were conserved, both Y450A-on and Y450A-mm6 showed elevated free-energy barriers at phase H (TS10), with TS10-OT = 11.27 kcal/mol and TS10-mm = 14.29 kcal/mol, respectively, compared to 7.02 kcal/mol in wt-on. In the Y450Amm6 system, the combined effects of Y450 removal and base mismatch further impaired DNA bending, leading to a significant increase in the energy barrier at TS9-mm (17.71 kcal/mol), which occurred even before maximum bending was achieved (vs. 8.54 kcal/mol in wild-type). These results demonstrate that Y450-tDNA interactions not only stabilize bent DNA conformations and promote accurate seed pairing, but also inadvertently stabilize mismatched pairs—particularly at the 6th position—thereby increasing off-target effects.

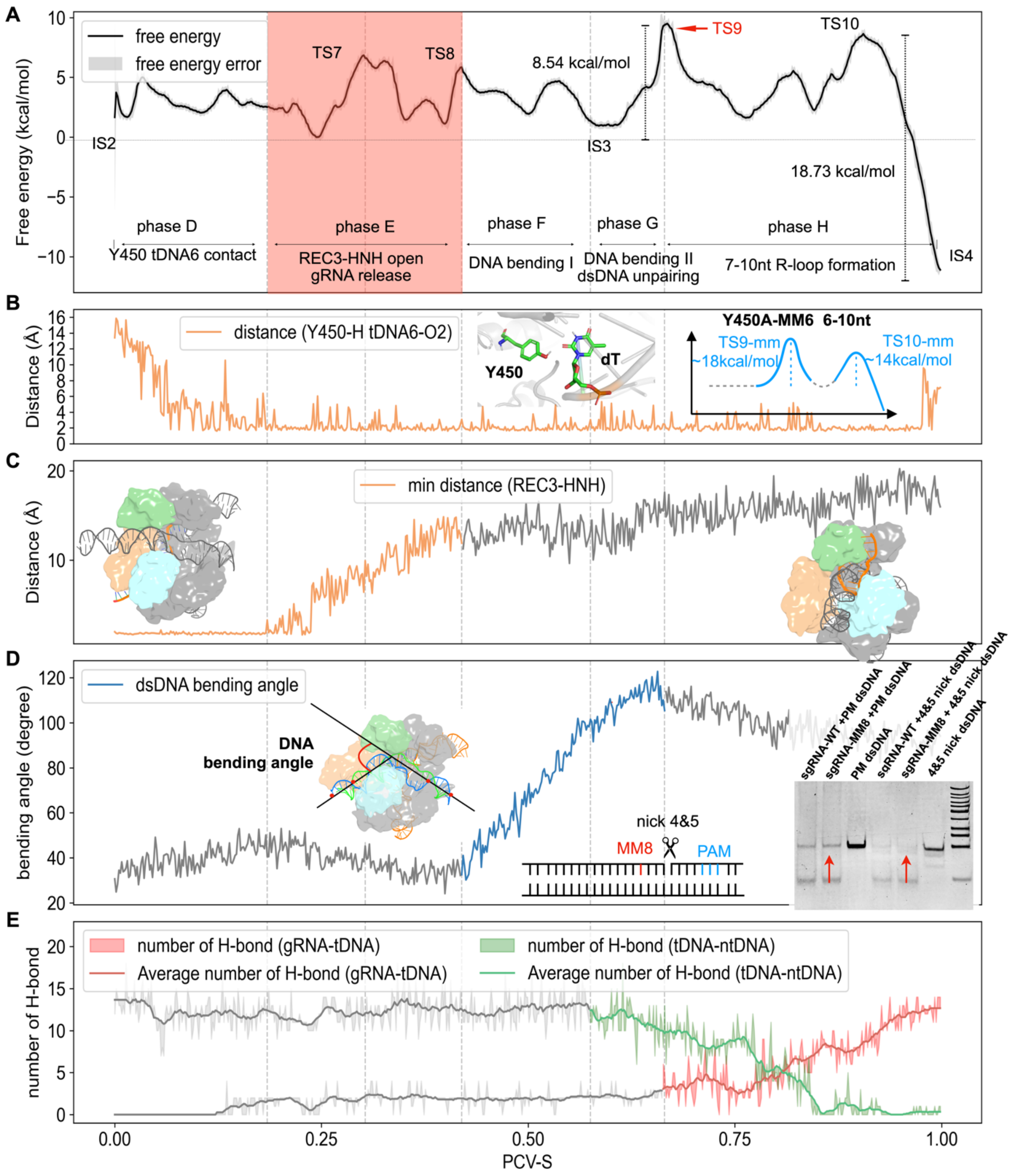


**Supplementary Fig. 3 | Kinetic mechanism of target recognition at 6-10 nt in the seed region of SpyCas9. a**, Free-energy profile for 6-10 nt recognition. **b**, Distance between Y450 and the 6th tDNA position, highlighting the stabilizing interaction between Y450 and the tDNA nucleobase during DNA bending. The energy schematic "Y450A-mm6" denotes the Y450A mutation with a gRNA-tDNA mismatch at position 6, and in this diagram the energy barriers for TS9-mm6 (~18 kcal/mol) and TS10-mm6 (~14 kcal/mol) are both markedly elevated compared to the wild-type on-target TS9 (~8.5 kcal/mol) and TS10 (~7 kcal/mol) in the 6-10nt region. **c**, Minimum distance between REC3 and HNH, reflecting the separation of the HNH–REC3 interface and the formation of a transient gap that permits gRNA release. **d**, DNA bending angle during the pairing process, showing the highest bending angle at transition state 9 (TS9, free-energy barrier ~8.54 kcal/mol). **e**, Changes in hydrogen bond numbers between gRNA-tDNA and ntDNA-tDNA. In phase G, DNA begins to unwind; in phase H, new gRNA-tDNA pairing initiates.

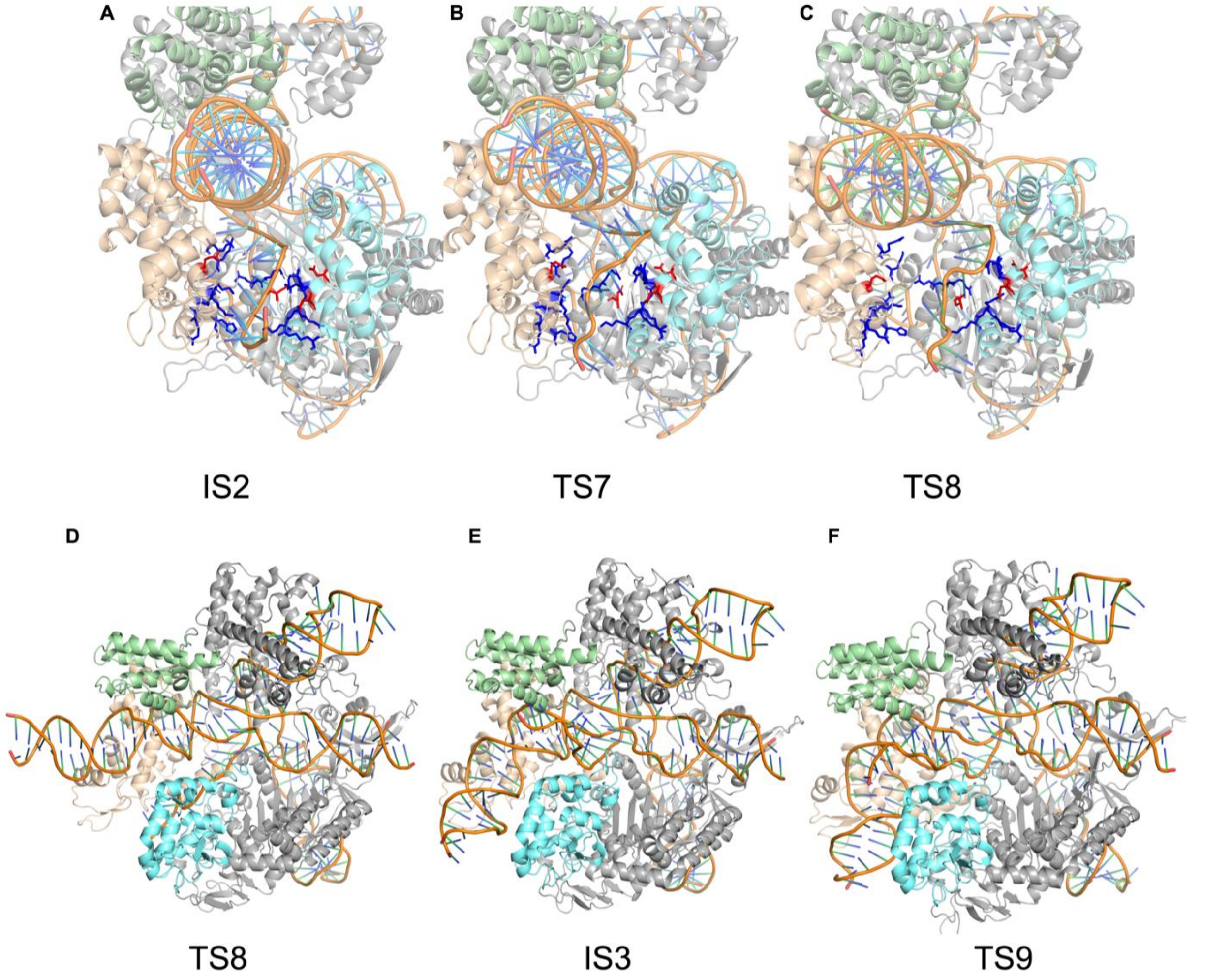


**Supplementary Fig. 4 | Structural snapshots of IS2, TS7, TS8, IS3 and TS9 during the 6-10 nt pairing process of CRISPR-SpyCas9.** As the gap between Rec3 and HNH opens, the previously buried 3′ end of the gRNA gradually overcomes the electrostatic barrier created by positive charged residues (blue)and is released into the solvent, facilitating the downstream base-pairing steps. Rec2 domain colored green, Rec3 domain colored wheat, HNH domain colored cyan.

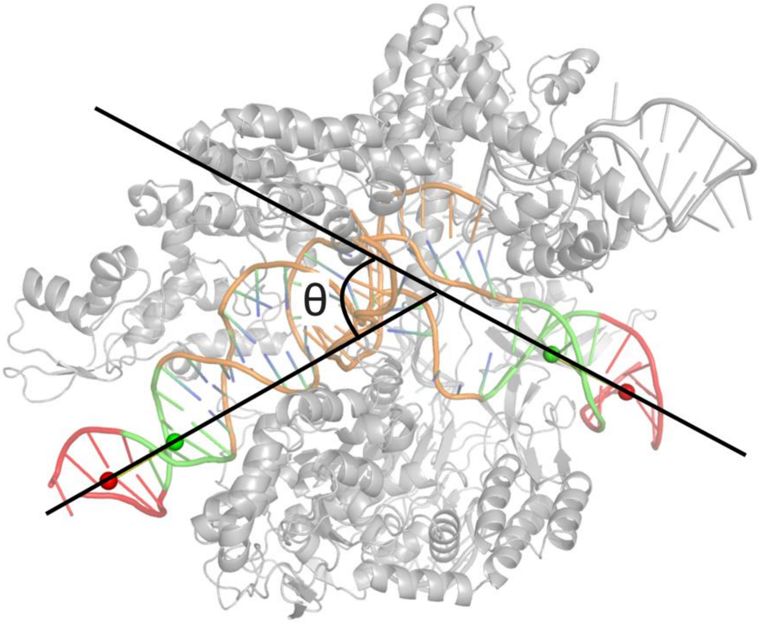


**Supplementary Fig. 5 | Calculation of DNA bending angle**. gRNA in orange. Red and green mark dsDNA segments on the left and right sides used for centroid calculation. For each side, compute the centroids of the red and green segments and define the side vector as the line from the green centroid to the red centroid:

$$v_L = C_L^{\text{red}} - C_L^{\text{green}}, \qquad v_R = C_R^{\text{red}} - C_R^{\text{green}}.$$

$$\theta = \arccos\left(\frac{|v_L \cdot v_R|}{|v_L|\,|v_R|}\right)$$

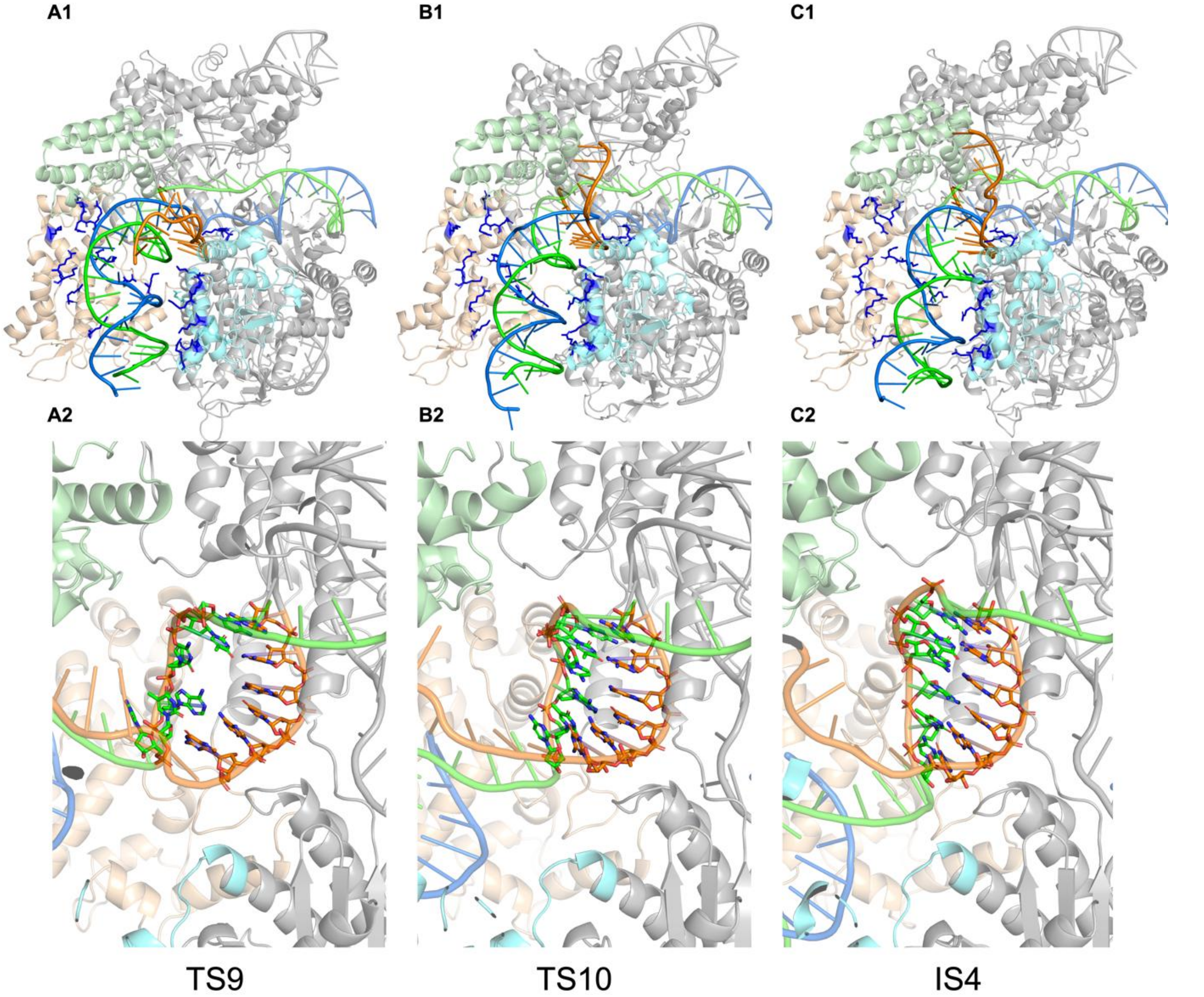


**Supplementary Fig. 6 | Structural snapshots of CRISPR-SpyCas9. Panels A1-B1-C1.** Structural snapshots of TS9, TS10, and the intermediate state (IS4) showing the distal (PAM-proximal) DNA after bending. As the DNA duplex bends, electrostatic interactions between the DNA backbone and positively charged residues lining the Rec3-HNH cleft become increasingly pronounced, while the minor groove at the PAM-distal end progressively widens to accommodate gRNA invasion. **Panels A2-B2-C2.** Corresponding snapshots of the 6-10 nt pairing process in TS9, TS10, and IS, with tDNA colored green and gRNA colored orange. These panels illustrate the stepwise expansion of the RNA-DNA hybrid: initial pairing in TS9, further maturation in TS10, and stabilization in the intermediate state 4.

In the Phase H (TS9→TS10), the PAM-distal segment of the duplex is progressively threaded into the positively charged REC3-HNH cavity (residues 767-769, 771-772, 774-778, 782-783, 789, 820, 877-878, 880-881, 884-885, 888, 890, 893-896, 899, 901-902, 905, 909, 918-919, 924-926, Supplementary Fig. 6 A1-C1). Electrostatic contacts with these basic residues tug on both tDNA and ntDNA, bending the duplex and prying the strands apart as the remaining 6-10 nt base pairs are sequentially disrupted. After hydrogen bonds in the 6-10 nt DNA duplex is fully disrupted, the system traverses TS10. Beyond TS10, the PAM-distal DNA continues to reposition within the cavity. Under the attractive field of basic residues on the HNH surface, tDNA and ntDNA are further splayed, opening a groove that allows the gRNA 3′ end to invade and establish a stable 6-10 nt hetero-duplex. This relaxation lowers the free energy by ~18.73 kcal $mol^{-1}$ and defines the stable intermediate IS4, signifying completion of seed recognition (Supplementary Fig. 6 A2-C2).

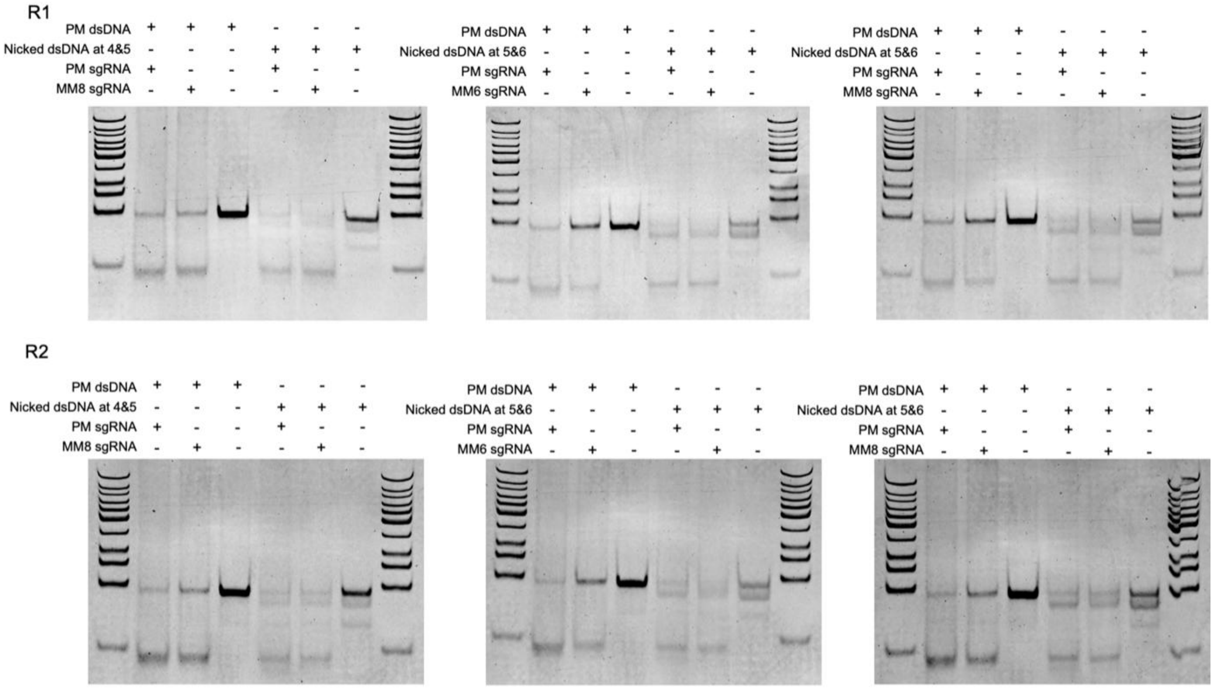


**Supplementary Fig. 7 | The results of Cas9 cleavage with dsDNA substrates.** PM dsDNA, perfect matched dsDNA; Nicked dsDNA at 4&5, dsDNA with a nick at position 4 and 5 from PAM; Nicked dsDNA at 5&6, dsDNA with a nick at position 5 and 6 from PAM; PM sgRNA, a sgRNA perfectly matched to PM dsDNA; MM8 sgRNA, a sgRNA mismatched to PM dsDNA at position 8 from PAM; MM6 sgRNA, a sgRNA mismatched to PM dsDNA at position 6 from PAM. The experiments were performed twice.

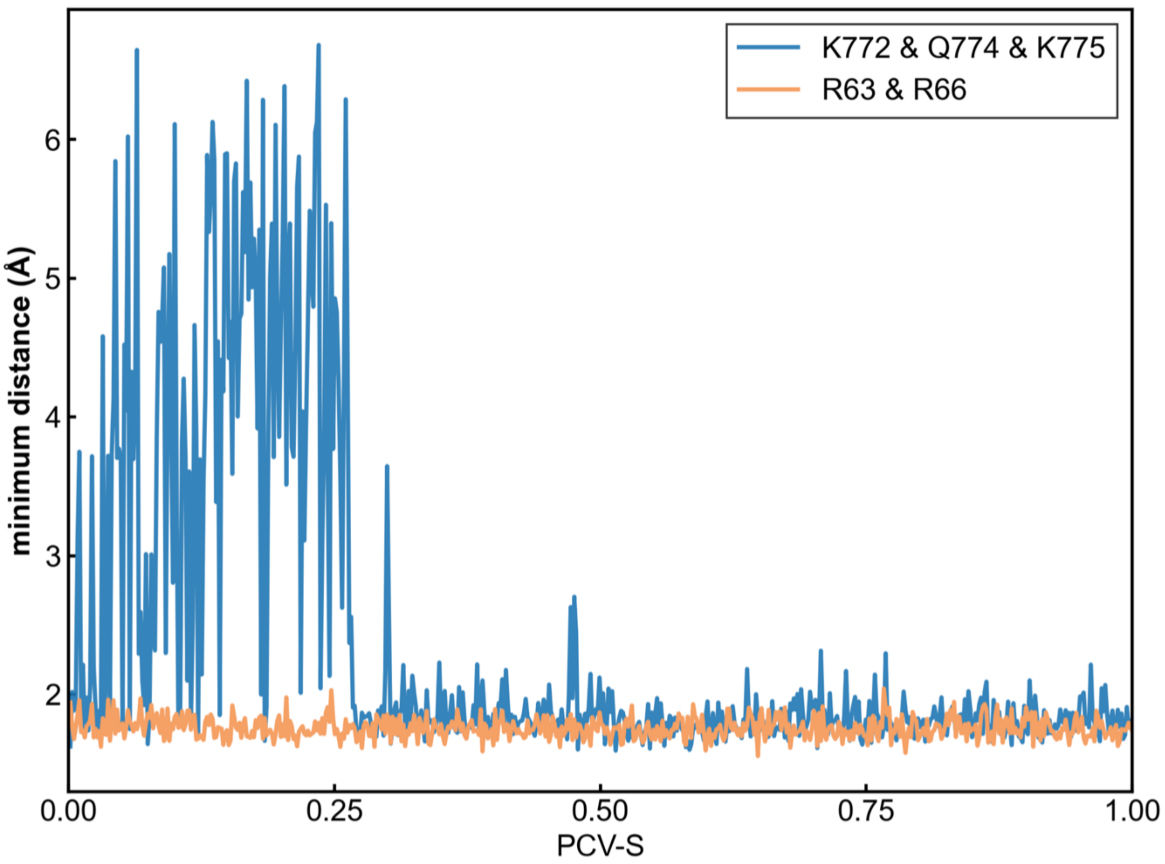


**Supplementary Fig. 8 | The minimum distance between key protein residues and gRNA during the 6-10 nt base-pairing process of CRISPR-Cas9.** The blue trajectory represents the minimum distance from the R63/R66 cluster to the gRNA, while the orange trajectory corresponds to the K772/Q774/K775 cluster.

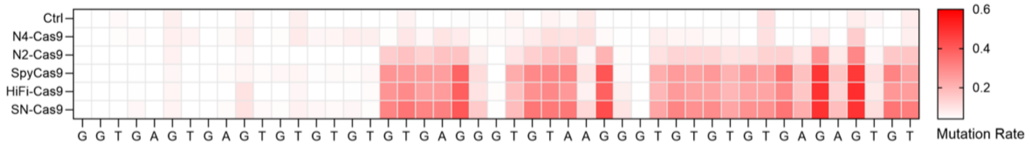


**Supplementary Fig. 9 | Base-resolution mutation rate profile at the OT1 site for different Cas9 variants.** Mutation rate = Heterozygous peaks value/Total peaks value.

| | Indel | Percentage | On-target site Sequence | |
|---|---|---|---|---|
| SpyCas9 | WT | 23% | TGTG**GGTGAGTGAGTGTGTGC** | **GTGTGG**GGTTGAGGGTG |
|  | -6 | 2% | TGTGGGTGAGTGAGTGTGTG- | -----GGGTTGAGGGTG |
|  | -8 | 1% | TGTGGGTGAGTGAGTGTGTG- | -------GTTGAGGGTG |
| | +1 | 71% | TGTGGGTGAGTGAGTGTGTGC | nGTGTGGGGTTGAGGGT |
| | +2 | 1% | TGTGGGTGAGTGAGTGTGTGC | nnGTGTGGGGTTGAGGG |
| N4-Cas9 | WT | 24% | TGTG**GGTGAGTGAGTGTGTGC** | **GTGTGG**GGTTGAGGGTG |
|  | -6 | 5% | TGTGGGTGAGTGAGTGTGTG- | -----GGGTTGAGGGTG |
|  | -8 | 4% | TGTGGGTGAGTGAGTGTGTG- | -------GTTGAGGGTG |
|  | -8 | 1% | TGTGGGTGAGTGAGTGTGT-- | ------GGTTGAGGGTG |
| | +1 | 60% | TGTGGGTGAGTGAGTGTGTGC | nGTGTGGGGTTGAGGGT |
| | +2 | 3% | TGTGGGTGAGTGAGTGTGTGC | nnGTGTGGGGTTGAGGG |

| | Indel | Percentage | Off-target site 1 (OT1) Sequence | |
|---|---|---|---|---|
| SpyCas9 | WT | 78% | TGAG**GGTGAGTGAGTGTGTGT** | **GTGAGG**GTGTAAGGGTG |
| | +1 | 20% | TGAGGGTGAGTGAGTGTGTGT | nGTGAGGGTGTAAGGGT |
| N4-Cas9 | WT | 100% | TGAG**GGTGAGTGAGTGTGTGT** | **GTGAGG**GTGTAAGGGTG |

| | Indel | Percentage | Off-target site 2 (OT2) Sequence | |
|---|---|---|---|---|
| SpyCas9 | WT | 72% | AGTG**GGTGAGTGAGTGCGTGC** | **GGGTGG**CGATGCAAGCG |
| | -4 | 5% | AGTGGGTGAGTGAGTGCG--- | -GGTGGCGATGCAAGCG |
|  | -8 | 2% | AGTGGGTGAGTGAGTGCG--- | -----GCGATGCAAGCG |
|  | -18 | 1% | AGTGGGTGAGTGAGTGCG--- | ---------------CG |
|  | -8 | 1% | AGTGGGTGAGTGAGTG----- | ---TGGCGATGCAAGCG |
| | +1 | 17% | AGTGGGTGAGTGAGTGCGTGC | nGGGTGGCGATGCAAGC |
| N4-Cas9 | WT | 98% | AGTGGGTGAGTGAGTGCGTGC | GGGTGGCGATGCAAGCG |
| | +1 | 1% | AGTGGGTGAGTGAGTGCGTGC | nGGGTGGCGATGCAAGC |

| | Indel | Percentage | Off-target site 3 (OT3) Sequence | |
|---|---|---|---|---|
| SpyCas9 | WT | 55% | TAAC**GCTGAGTGAGTGTATGC** | **GTGTGG**CTTTAGCGGGA |
|  | -8 | 6% | TAACGCTGAGTGAGTGTATGC | --------TTAGCGGGA |
|  | -12 | 4% | TAACGCTGAGTGAGTGTATGC | ------------CGGGA |
| | +1 | 31% | TAACGCTGAGTGAGTGTATGC | nGTGTGGCTTTAGCGGG |
| N4-Cas9 | WT | 82% | TAACGCTGAGTGAGTGTATGC | GTGTGGCTTTAGCGGGA |
|  | -8 | 1% | TAACGCTGAGTGAGTGTATGC | --------TTAGCGGGA |
| | +1 | 16% | TAACGCTGAGTGAGTGTATGC | nGTGTGGCTTTAGCGGG |

**Supplementary Fig. 10 | Detailed analysis of insertion and deletion (indel) profiles generated by SpyCas9 and N4-Cas9 at on-target and off-target sites (OT1, OT2, and OT3).**

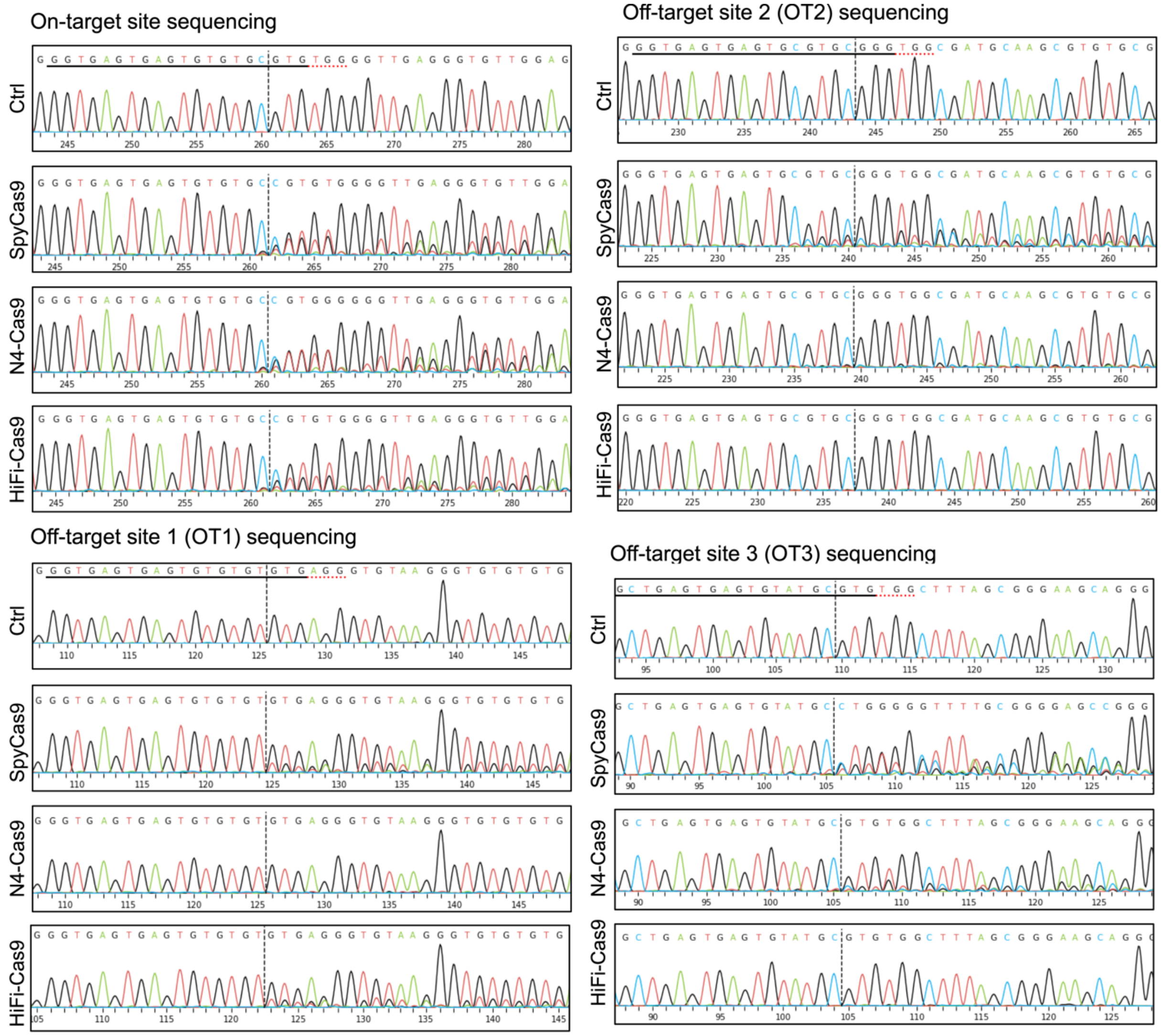


**Supplementary Fig. 11 | Representative sanger sequencing chromatograms for the OT1, OT2 and OT3 sites for different Cas9 variants.**

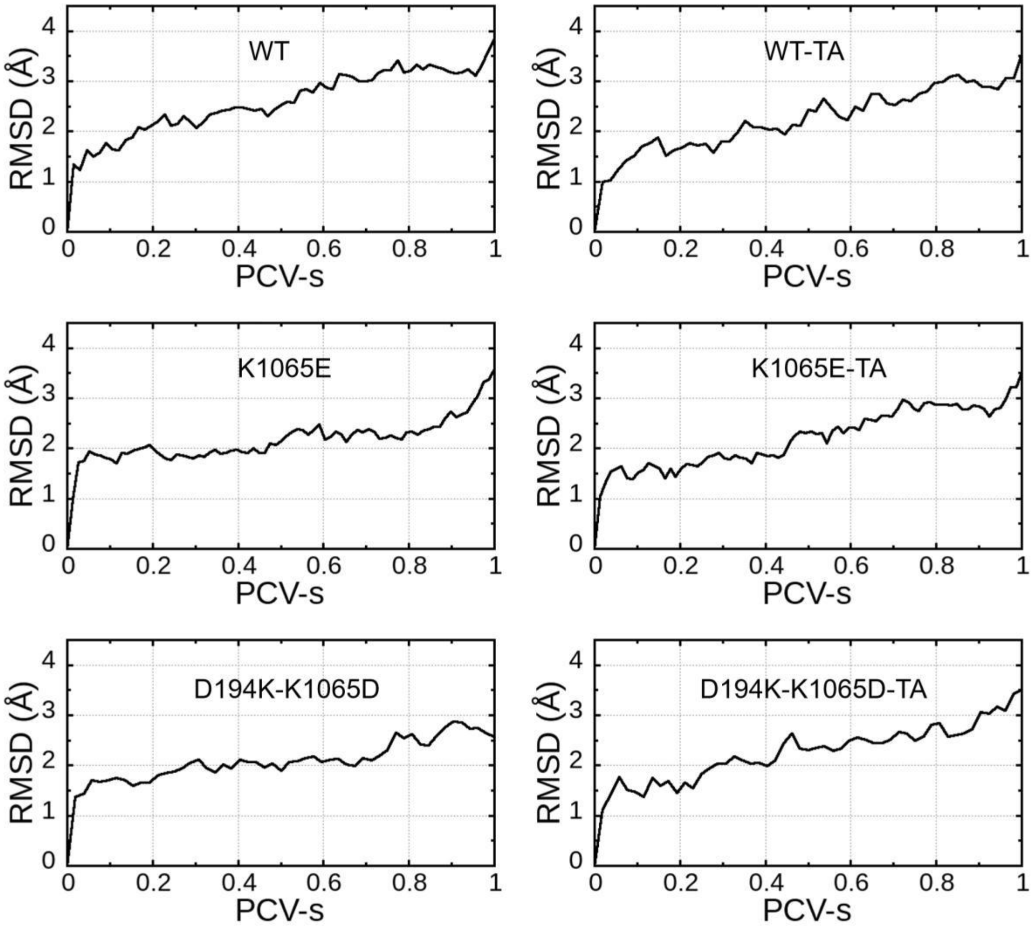


**Supplementary Fig. 12 | RMSDs of protein CA atoms for six paths of Cas12a.** No large protein conformation changes were observed during the base-pairing process of seed region.

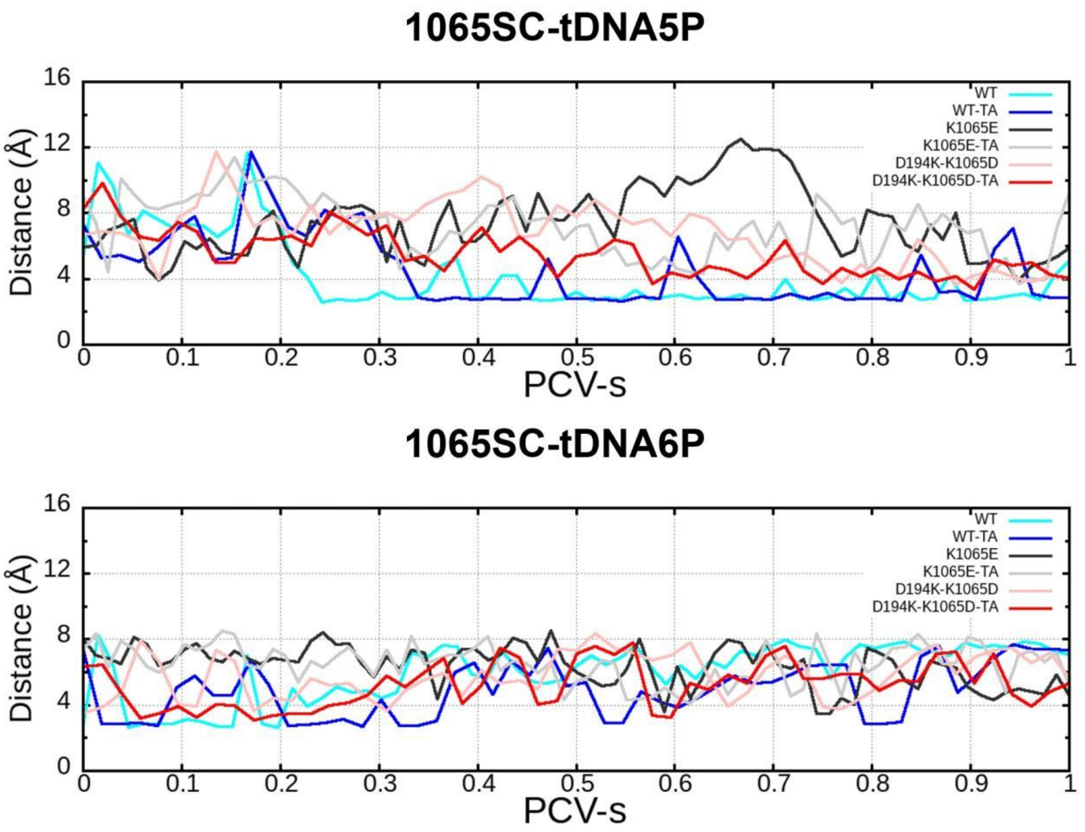


**Supplementary Fig. 13 | Distances between 1065 sidechain (SC) and tDNA5/6 phosphate (P) in the basing pairing processes of CRISPR-Cas12a.**

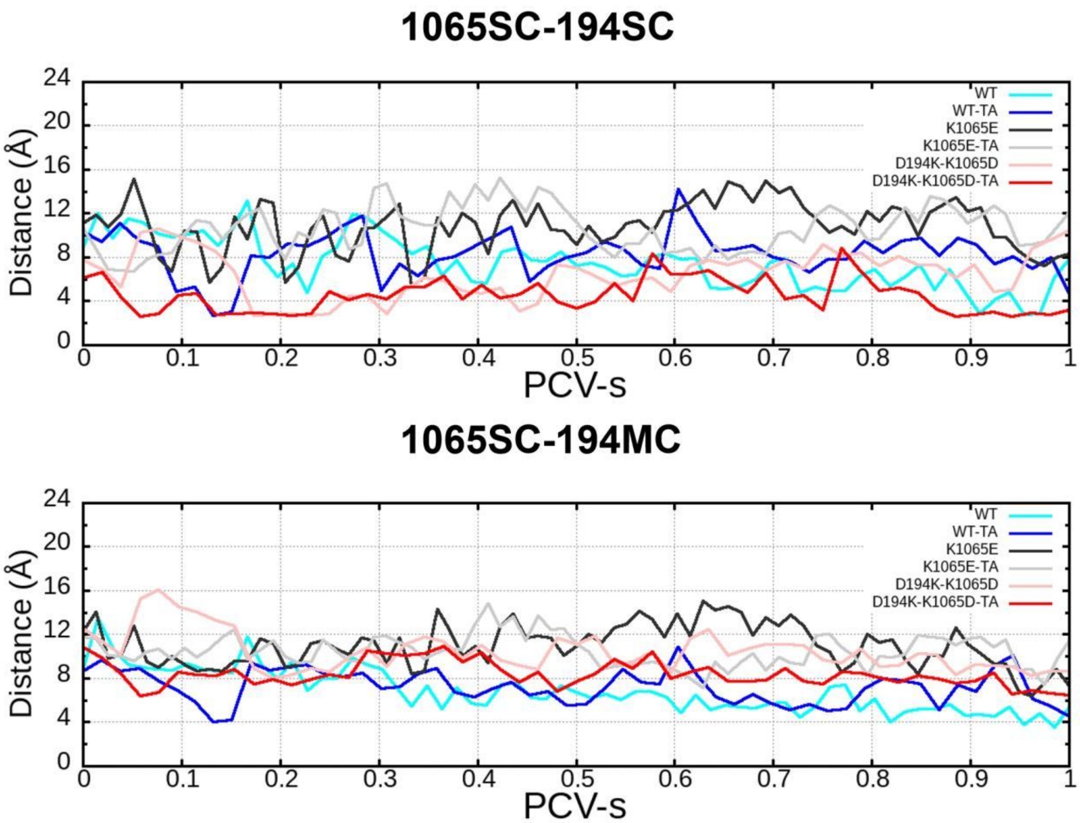


**Supplementary Fig. 14 | Distances between 1065 sidechain (SC) and D194 SC/MC (mainchain O) in the basing pairing processes of CRISPR-Cas12a.**

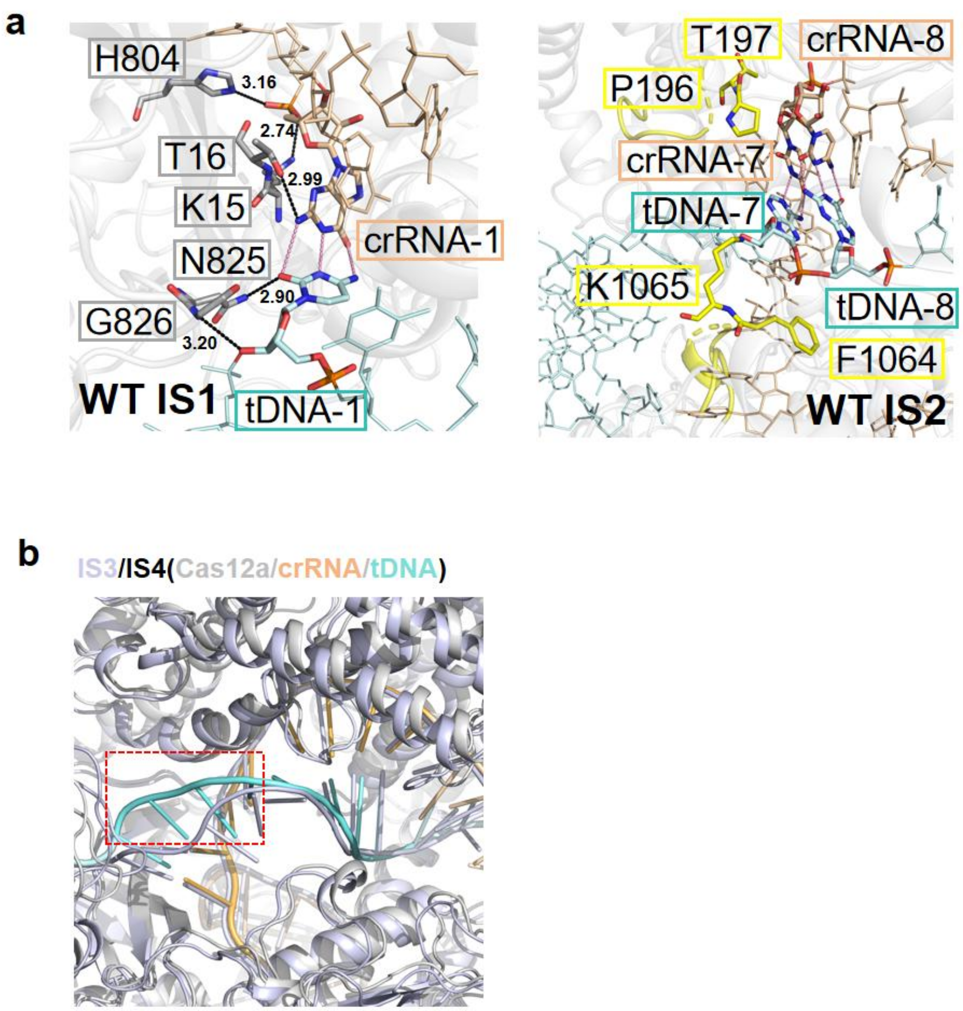


**Supplementary Fig. 15 | Intermediate states structures of WT Cas12a with perfect match DNA.**

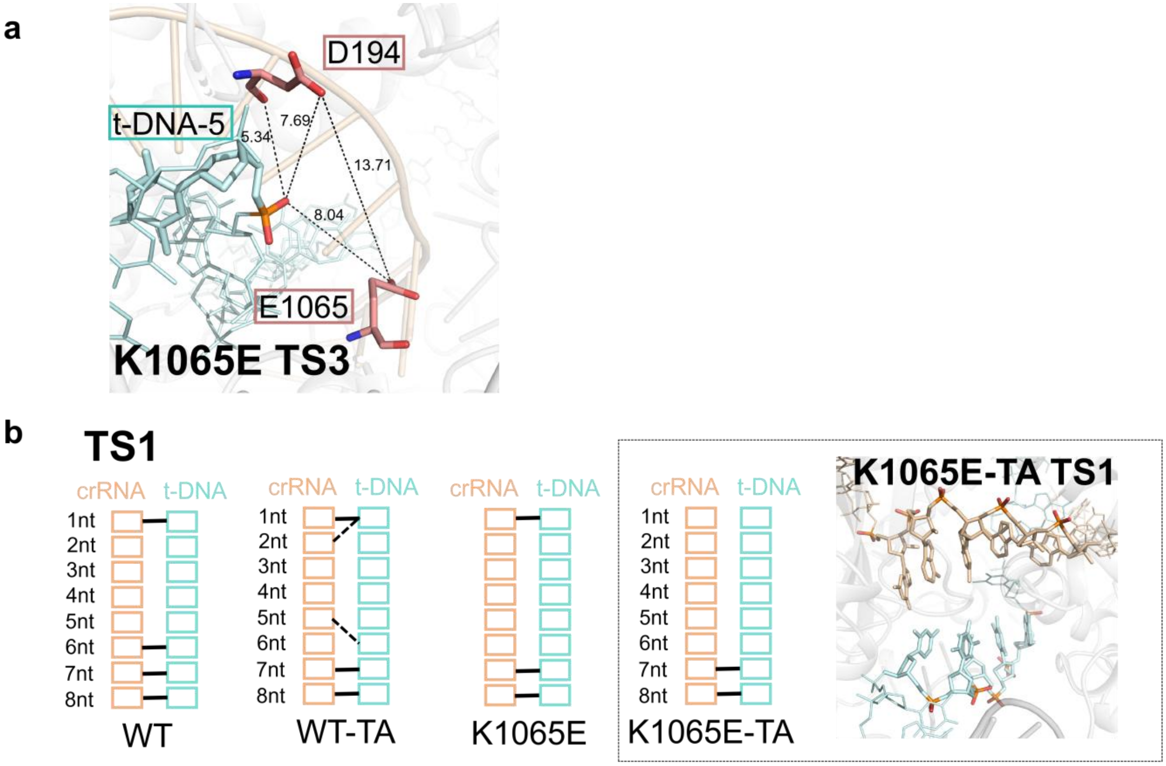


**Supplementary Fig. 16 | Transition states of Cas12a K1065E mutant with perfect match DNA (a) and TA mismatched DNA (b).**

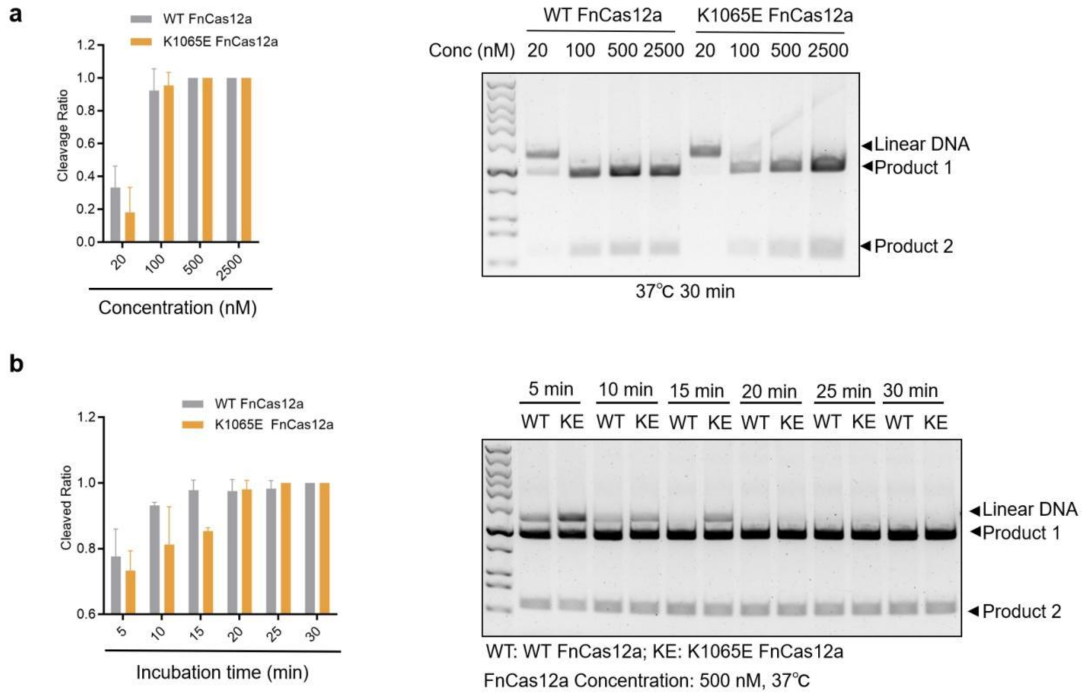


**Supplementary Fig. 17 | Optimization of experiment condition for FnCas12a.**

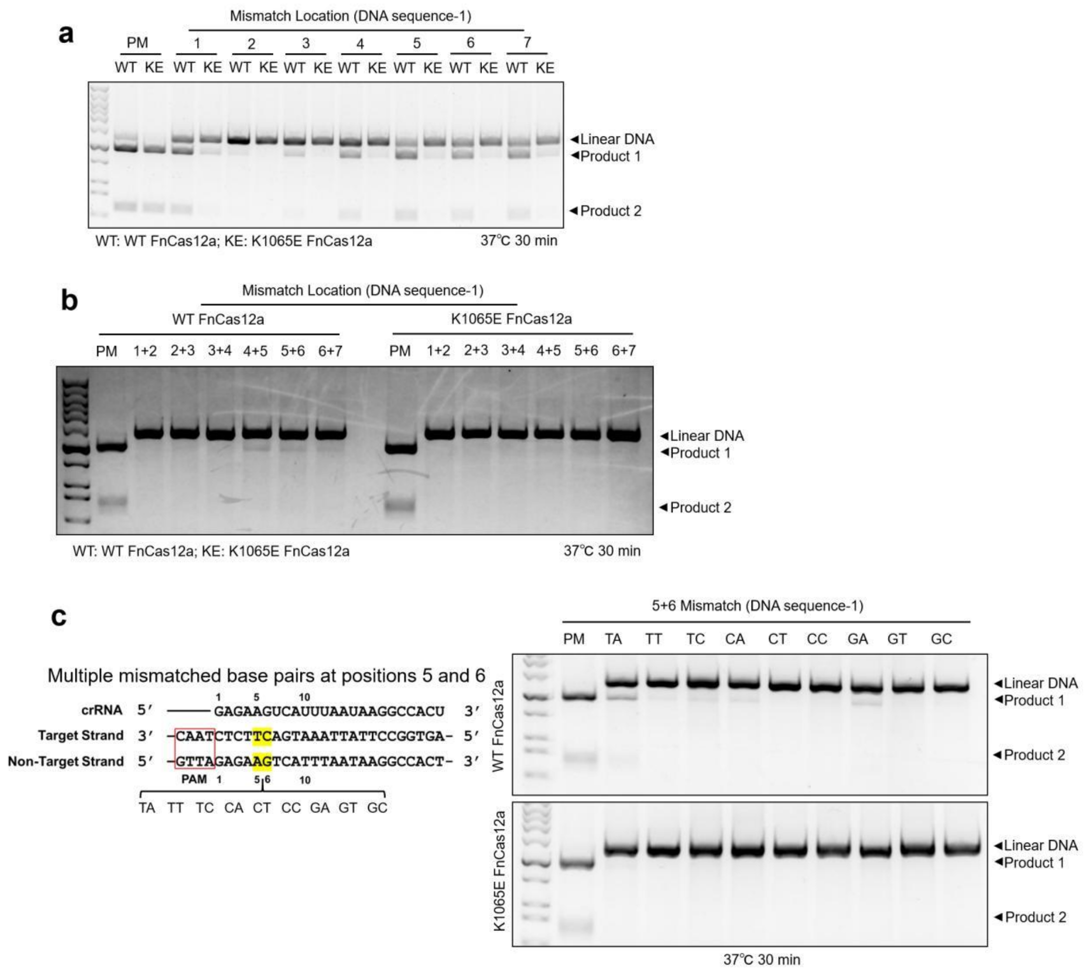


**Supplementary Fig. 18 | Experimental results for cleavage of sequence 1 by FnCas12a WT and K1065E mutant.** One of three replicas is shown for (**a-c**) *cis*-cleavage respectively.

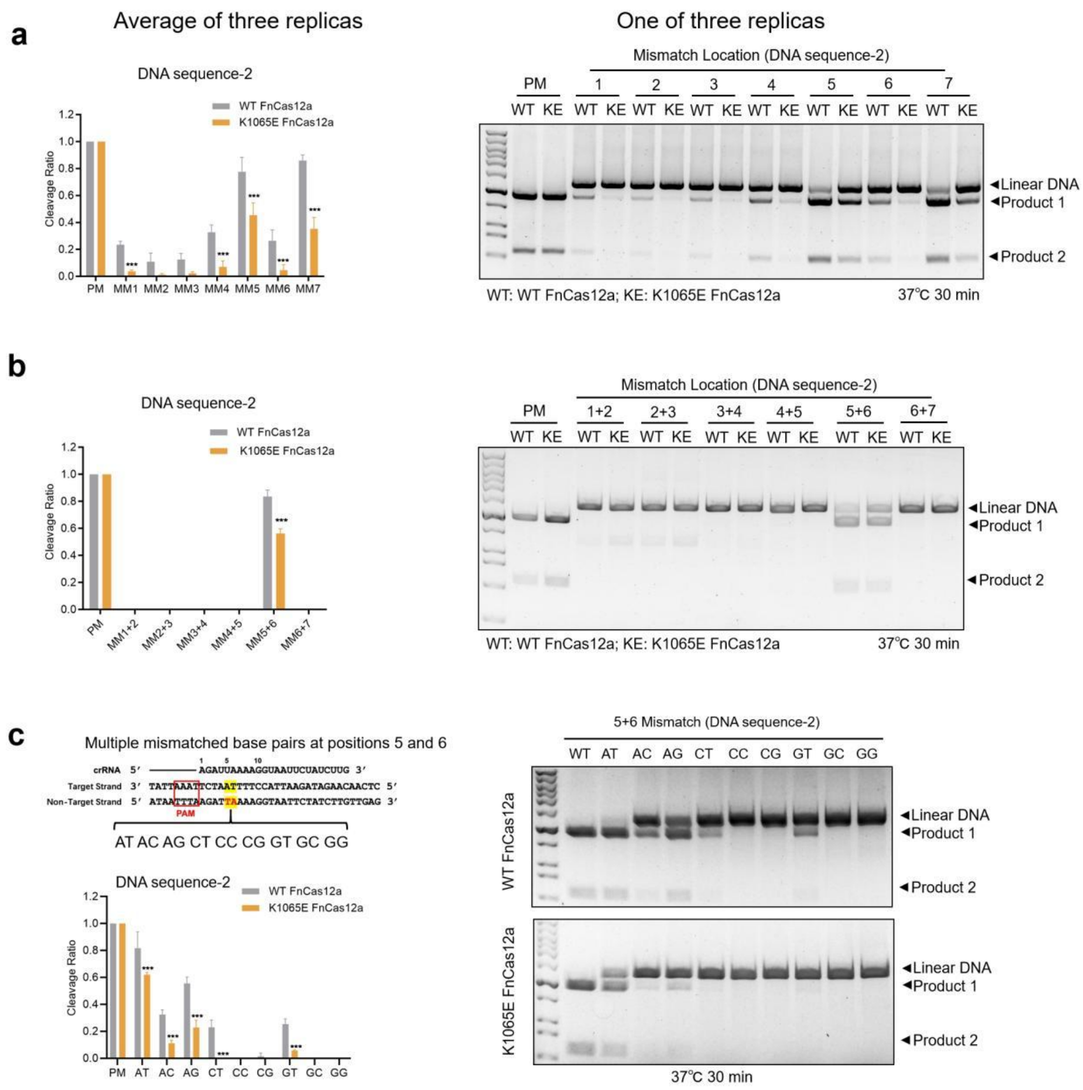


**Supplementary Fig. 19 | Experimental results for *cis*-cleavage of sequence 2 by FnCas12a WT and K1065E mutant.**

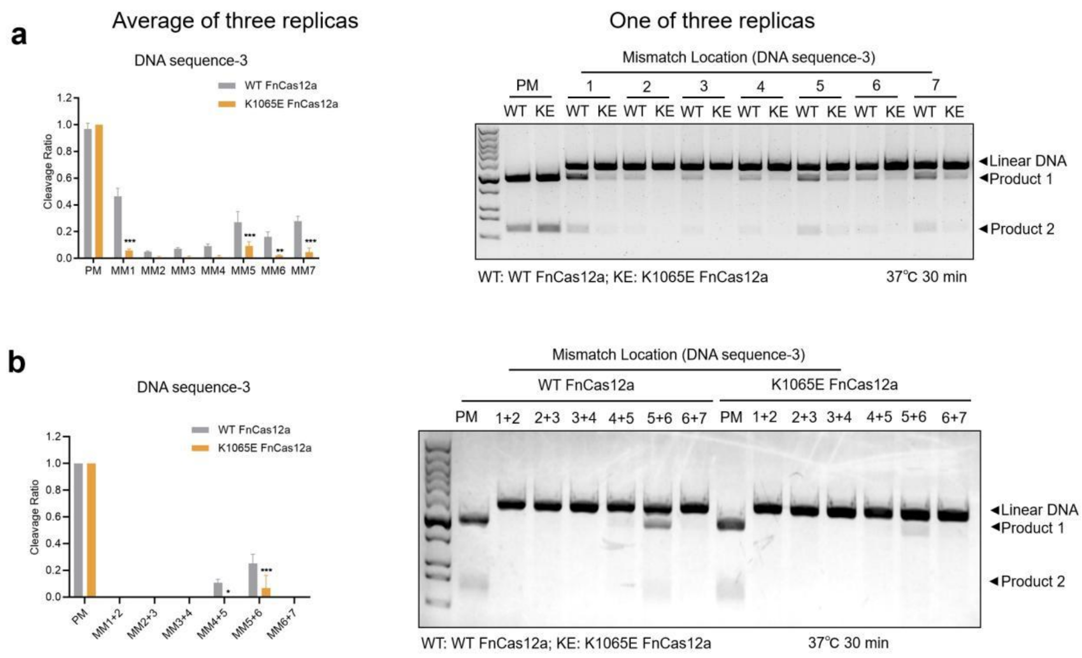


**Supplementary Fig. 20 | Experimental results for *cis*-cleavage of sequence 3 by FnCas12a WT and K1065E mutant.**

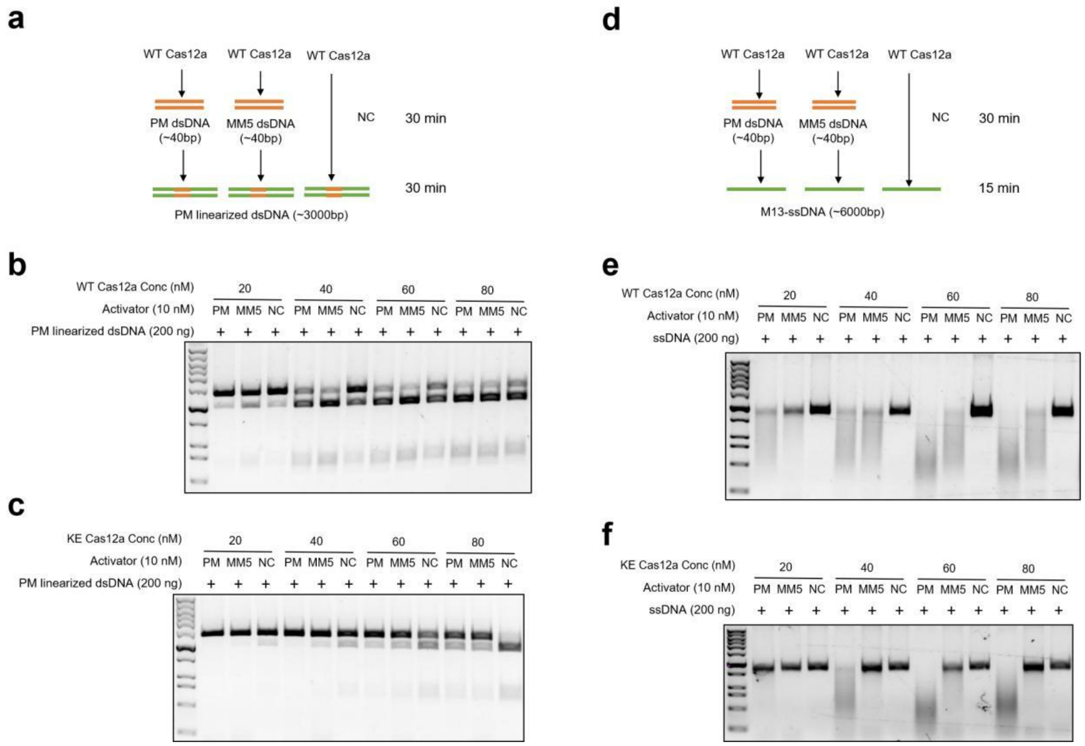


**Supplementary Fig. 21 | The *cis*- and *trans*-activity of sequence 1 for FnCas12a K1065E variant.**

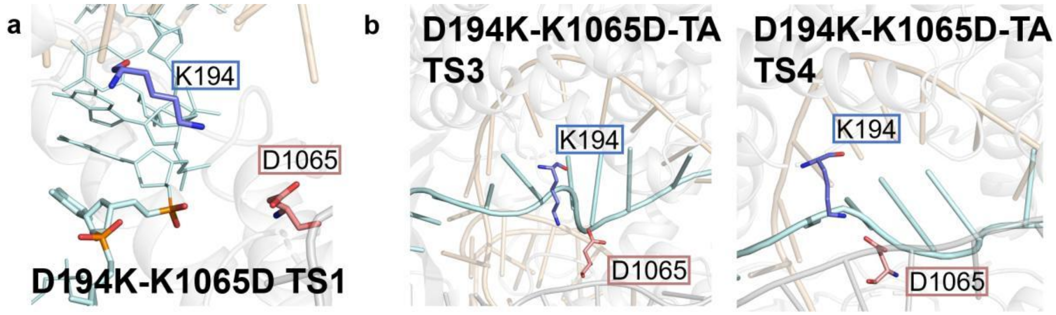


**Supplementary Fig. 22 | Transition states of CRISPR-FnCas12a D194K-K1065D mutant simulations.**

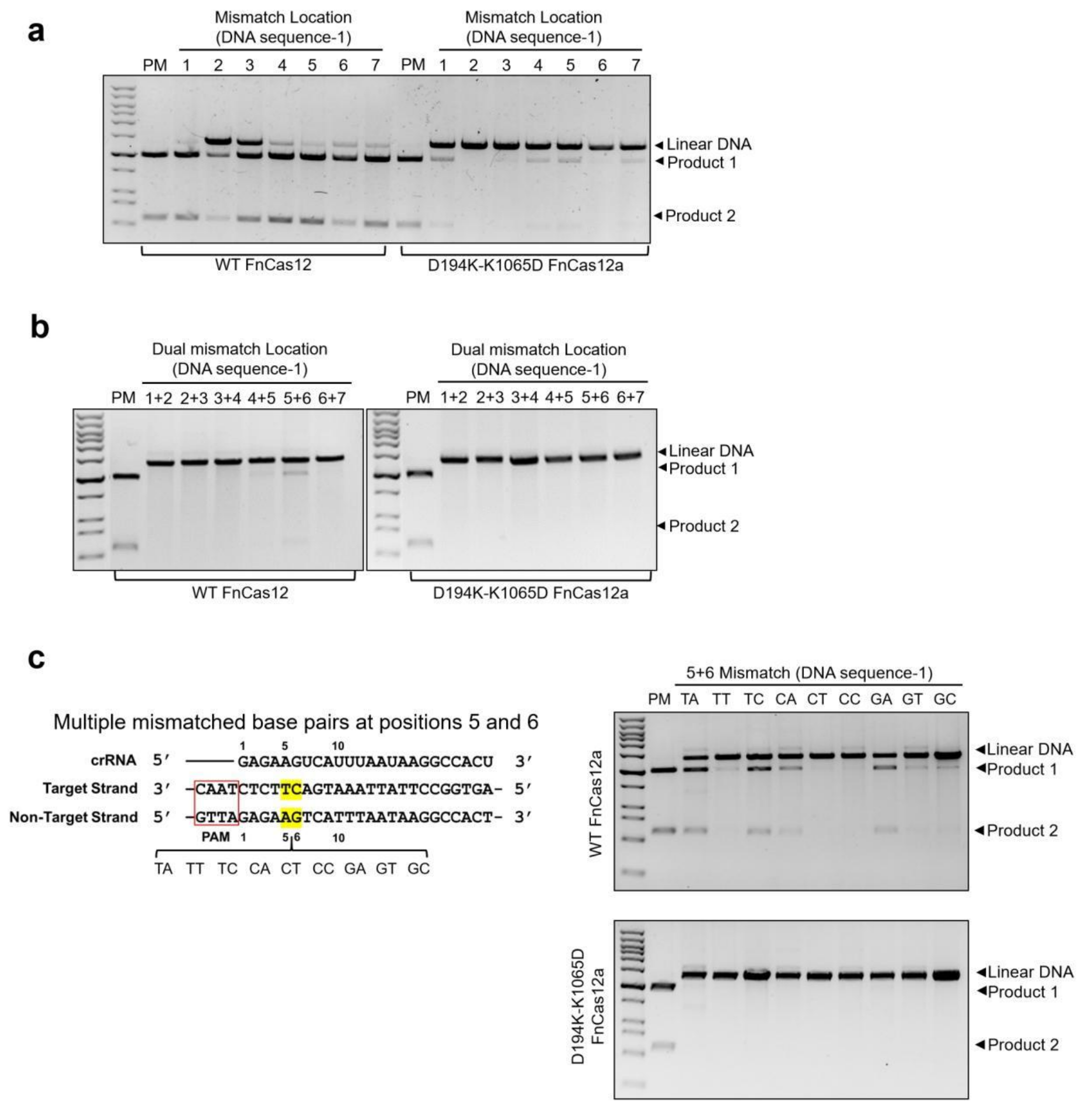


**Supplementary Fig. 23 | Experimental results for cleavage of sequence 1 by FnCas12a WT and D194K-K1065D mutant.** One of three replicas is shown for (**a-c**) *cis*-cleavage respectively.

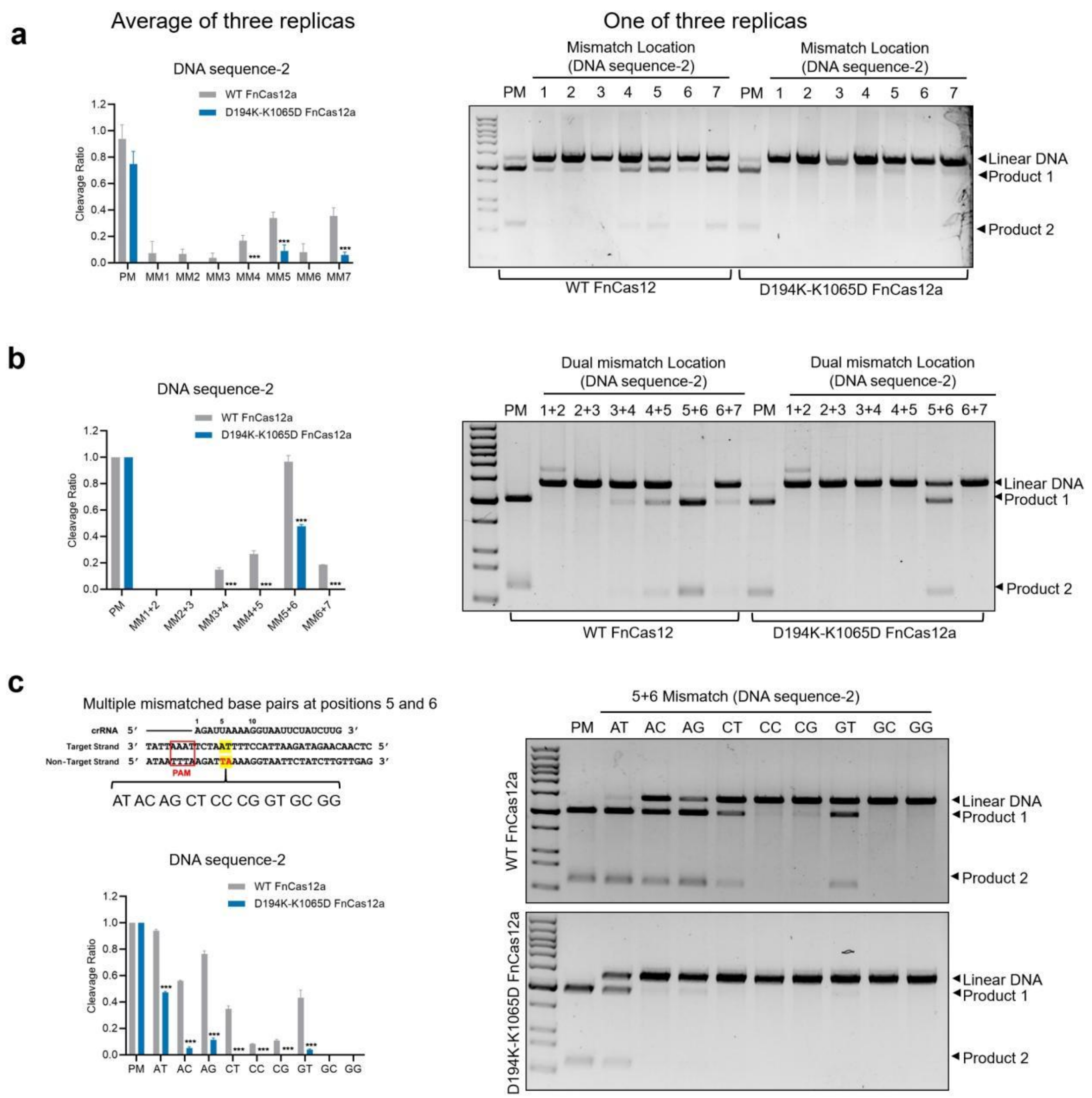


**Supplementary Fig. 24 | Experimental results for *cis*-cleavage of sequence 2 by FnCas12a WT and D194K-K1065D mutant.**

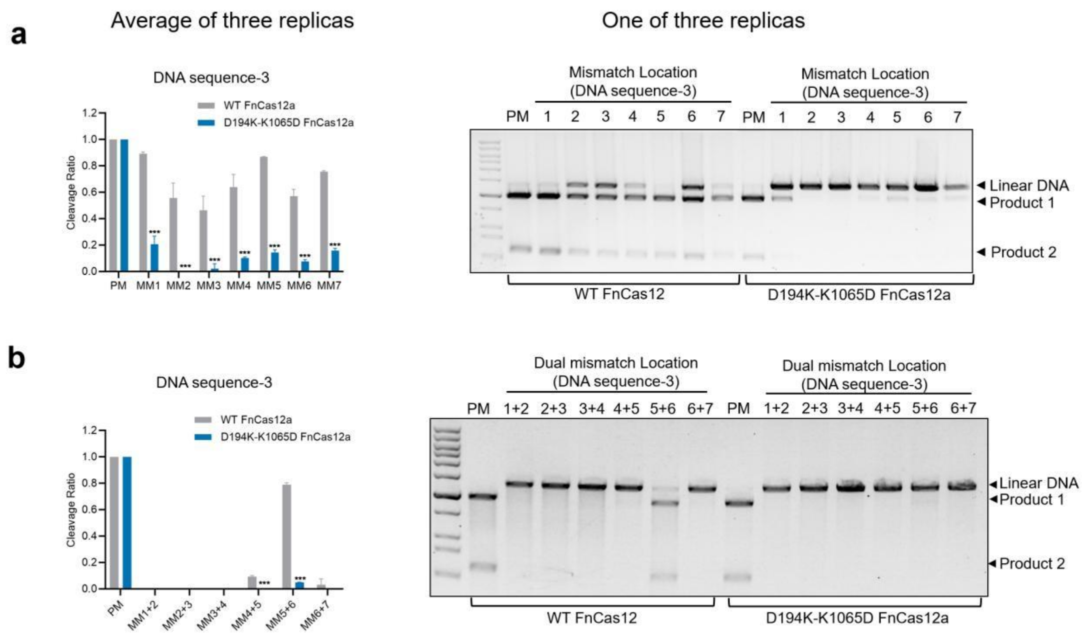


**Supplementary Fig. 25 | Experimental results for *cis*-cleavage of sequence 3 by FnCas12a WT and D194K-K1065D mutant.**

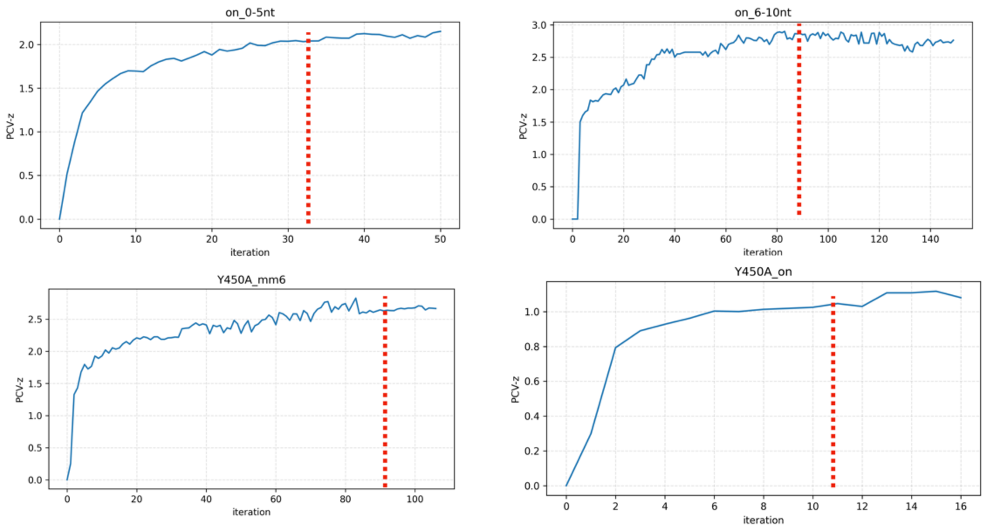


**Supplementary Fig. 26 | Convergence test by $\sqrt{\langle z \rangle}$ calculation for three paths of SpyCas9.**

Convergence of the optimization processes for the paths is measured by the progress of $\sqrt{\langle z \rangle}$ along the simulation iteration. The initial paths are the reference. Convergence iteration is highlighted by a red dashed line.

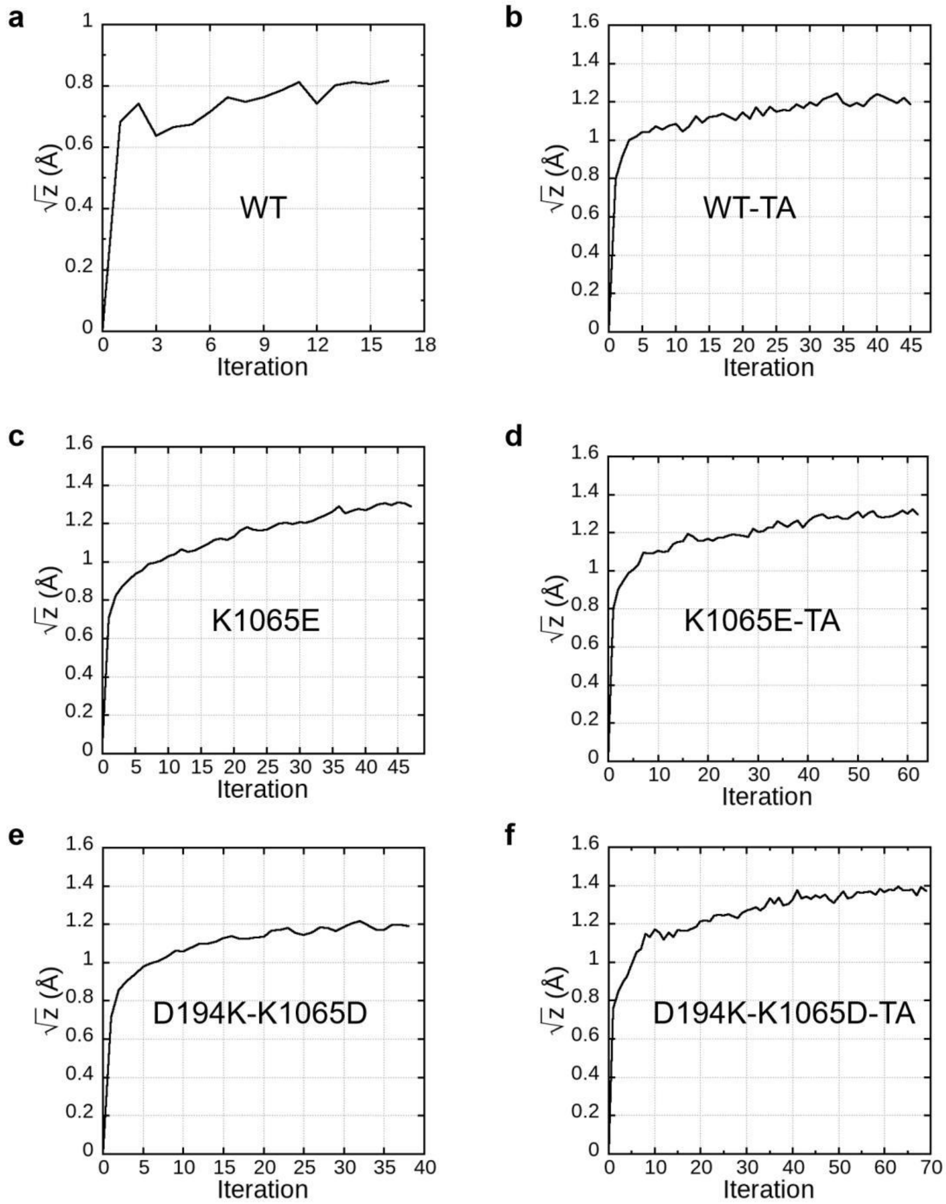


**Supplementary Fig. 27 | Convergence of TAPS simulations for CRISPR-Cas12a.**

**Supplementary Table 1 | Charged residues driving base pairing at positions 1-5 in Cas9 seed region.**

| Protein domain | | |
|---|---|---|
| Rec2 | negatively charged residues | E197, E198, D257, E260, D261, D269, D272-274, D275, D284, D288 |
| | positively charged residues | K218, R220, R221, K233, K234, K253, K263, K268 |
| Rec3 | positively charged residues | H511, R557, K558, K562, R586, H595, K599, K602, K604, K646, K649, K652, R653-655, R661 |

**Supplementary Table 2 | Nucleic acid sequences used in CRISPR-Cas12a MD.**

| Name | Sequences |
|---|---|
| crRNA(5’-3’) | AAUUUCUACUGUUGUAGAU-**GAGAAGUC**AUUUAAUAAGGC-CACU |
| PM-tDNA(3’-5’) | GCTCGAGCAAT-**CTCTTCAG**TAAATTATTCCG-GTGACGT |
| PM-ntDNA(5’-3’) | CGAGCTCGTTA-**GAGAAGTC**ATTTAATAAGGC-CACTGCA |
| DM-tDNA(3’-5’) | GCTCGAGCAAT-**CTCTATAG**TAAATTATTCCG-GTGACGT |
| DM-ntDNA(5’-3’) | CGAGCTCGTTA-**GAGATATC**ATTTAATAAGGC-CACTGCA |

The nucleic acids in sites 1-8 are labeled in bold and sites 5-6 are colored in red. PM: perfect match, DM: double mismatch.

**Supplementary Table 3 | Details of prepared Cas9 simulation system.**

| System | WT-OT 1-5nt | WT-OT 6-10nt | Y450A-OT | Y450A-MM6 |
|---|---|---|---|---|
| Main Components | Wt-spyCas9 + on-target sequence | Wt-spyCas9 + on-target sequence | Y450A-spyCas9 + on-target sequence | Y450A-spyCas9 + position 6 mismatch sequence ( dT-A) |
| TIP3 | 144157 | 144157 | 144157 | 144157 |
| $Cl^-$ | 724 | 724 | 724 | 724 |
| $K^+$ | 839 | 839 | 839 | 839 |
| $Mg^{2+}$ | 14 | 14 | 14 | 14 |
| Total Atoms | 461839 | 461839 | 461828 | 461829 |
| Temperature | 310k | 310k | 310k | 310k |
| Pressure | 1 atm | 1 atm | 1 atm | 1 atm |

**Supplementary Table 4. Details of tMD simulations for Cas9.**

| Initial Path | | WT-OT 1-5nt | WT-OT 6-10nt |
|---|---|---|---|
| Initial State | | PDB ID: 7S36, equilibrated after 100 ns MD simulation | IS2 from path WT-OT 1-5nt |
| Target State | | 3 nt structure (PDB ID: 7S38)<br>5 nt structure (PDB ID: 7Z4C) | 8 nt structure (PDB ID: 7Z4E)<br>12 nt structure (PDB ID: 7Z4G) |
| Temperature | | 310k | 310k |
| Pressure | | 1 atm | 1 atm |
| Atom Sets | Alignment | Cα of helical part in RuvC-1, BH,REC1,WED and PI domain | |
| | RMSD | Heavy atoms of target strand DNA 1-5nt and REC2, REC3 HNH domain | Heavy atoms of target strand DNA 1-5nt, sgRNA5-10nt and REC2,REC3 HNH domain |
| Force constant | | 0 to 200,000 kJ/mol/nm2 | 0 to 50,000 kJ/mol/nm2 |
| Frame Record Frequence | | 10 ps | 10 ps |
| Total Sampling Time | | 10 ns | 12 ns |

**Supplementary Table 5 | Details of TAPS for Cas9.**

| Optimized Path | | WT-OT 1-5nt | WT-OT 6-10nt | Y450A-OT | Y450A-MM6 |
|---|---|---|---|---|---|
| Sampling Time of Each Iteration | | 4000 ps | | | |
| Temperature | | 310 K | | | |
| Pressure | | 1 atm | | | |
| Atoms Set | Alignment | Cα of helical part in RuvC-1, BH, REC1, WED and PI domain | | | |
| | RMSD | Heavy atoms of dsDNA, sgRNA 1-5nt and REC2, REC3 HNH domain | Heavy atoms of dsDNA, sgRNA 5-10nt and REC2, REC3 HNH domain | | |
| Tolerant Distance for Neighbor Nodes | | 1.15 Å | | | |
| Well-tempered Metadynamics simulation | Gaussian Height | 0.25 kJ/mol | | | |
| | Gaussian Width | 0.50 | | | |
| | Bias Factor | 10 | | | |
| Length of tMD | | 10 ps | | | |
| Force Constant of tMD | | 150,000 kJ/mol/nm2 | | | |
| Frame Record Frequence | | 1 ps | | | |

**Supplementary Table 6 | Details of Umbrella Sampling for Cas9.**

| Optimized Path | | WT-OT 1-5nt | WT-OT 6-10nt | Y450A-OT | Y450A-MM6 |
|---|---|---|---|---|---|
| Sampling time of each node | | 4 ns | | | |
| Temperature | | 310 K | | | |
| Pressure | | 1 atom | | | |
| Node Number in Final Path | | 62 | 68 | 80 | 91 |
| Window size | | 0.25 PCV-s | | | |
| Total Number of Windows for Umbrella Sampling | | 248 | 272 | 320 | 364 |
| PCV-based Umbrella Sampling | Position for z-Wall Potential | 0.0256 $nm^2$ | | | |
| | Force Constant for PCV-z | 3125000.0 $kJ/mol/nm^2$ | | | |
| | Force Constant for PCV-s | 600 kJ/mol | | | |

**Supplementary Table 7 | Three sequences of crRNA used in Cas12a experiments.**

| Name | Sequences (5'-3') |
|---|---|
| crRNA-1 | AAUUUCUACUGUUGUAGAU**GAGAAGUC**AUUUAAUAAGGCCACU |
| crRNA-2 | AAUUUCUACUGUUGUAGAU**AGAUUAAA**AGGUAAUUCUAUCUUG |
| crRNA-3 | AAUUUCUACUGUUGUAGAU**GUGAUAAG**UGGAAUGCCAUGUGGG |

The nucleic acids in sites 1-8 are labeled in bold. crRNA-1 is same to the sequence used in MD.

**Supplementary Table 8 | Zero-shot prediction of the most/second favorable mutations for Cas9 by protein language models (https://huggingface.co/spaces/thaidaev/zsp).**

| No. | Model | R63 (R63S rank) | R66 | K772 (K772S rank) | Q774 | K775 |
|---|---|---|---|---|---|---|
| 1 | esm1b_t33_650M_UR50S | R63R/K (5) | R66R/K | K772K/R (5) | Q774K | K775K/Q |
| 2 | esm1v_t33_650M_UR90S_1 | R63R/K (7) | R66R/E | K772K/E (7) | Q774L | K775K/E |
| 3 | esm1v_t33_650M_UR90S_5 | R63K (5) | R66K | K772E (3) | Q774L | K775K/E |
| 4 | esm2_t6_8M_UR50D | R63K (8) | R66K | K772K/E (3) | Q774K | K775K/L |
| 5 | esm2_t12_35M_UR50D | R63K (5) | R66K | K772K/R (3) | Q774S | K775K/R |
| 6 | esm2_t30_150M_UR50D | R63R/K (4) | R66R/K | K772R/Q (6) | Q774K | K775K/R |
| 7 | esm2_t33_650M_UR50D | R63R/K (6) | R66R/K | K772K/R (9) | Q774R | K775K/R |
| 8 | esm2_t36_3B_UR50D | R63R/K (8) | R66R/K | K772R (7) | Q774K | K775K/R |
| 9 | Profluent-Bio/E1-150m | R63R/G (3) | R66R/K | K772K/E (7) | Q774K | K775K/E |
| 10 | Profluent-Bio/E1-300m | R63R/S (2) | R66R/K | K772K/E (5) | Q774K | K775K/R |
| 11 | Profluent-Bio/E1-600m | R63R/T (5) | R66R/T | K772K/E (6) | Q774K | K775K/R |

**Supplementary Table 9 | Zero-shot prediction of the most/second favorable mutations for Cas12a by protein language models (https://huggingface.co/spaces/thaidaev/zsp).**

| No. | Model | D194 (D194K rank) | K1065 （K1065E rank） |
|---|---|---|---|
| 1 | esm1b_t33_650M_UR50S | D194K (1) | K1065K/T (7) |
| 2 | esm1v_t33_650M_UR90S_1 | D194K (1) | K1065K/L (4) |
| 3 | esm1v_t33_650M_UR90S_5 | D194K (1) | K1065I (4) |
| 4 | esm2_t6_8M_UR50D | D194K (1) | K1065K/E (2) |
| 5 | esm2_t12_35M_UR50D | D194K (1) | K1065K/E (2) |
| 6 | esm2_t30_150M_UR50D | D194K (1) | K1065E (1) |
| 7 | esm2_t33_650M_UR50D | D194D/E (7) | K1065K/E (2) |
| 8 | esm2_t36_3B_UR50D | D194E (7) | K1065K/Q (4) |
| 9 | Profluent-Bio/E1-150m | D194K (1) | K1065D (4) |
| 10 | Profluent-Bio/E1-300m | D194K (1) | K1065K/D (3) |
| 11 | Profluent-Bio/E1-600m | D194N (7) | K1065K/N (6) |
